# Basal BNIP3/NIX mitophagy is controlled by selective protection of sentinel receptors

**DOI:** 10.64898/2026.06.25.734517

**Authors:** Abhishek Kumar, Abigail C. Love, Keri-Lyn Kozul, Mehmet Oguz Gok, Natalie M. Niemi, Jonathan R. Friedman

## Abstract

Mitochondrial homeostasis is maintained by multiple quality control pathways, including mitophagy, which targets dysfunctional mitochondria for degradation. During receptor-mediated mitophagy, the outer membrane proteins BNIP3 and NIX directly recruit autophagy machinery to the mitochondrial surface, though their precise regulation is still unclear. In recent years, new BNIP3- and NIX-interacting proteins have been identified that influence mitophagic flux. PPTC7 and FBXL4 target BNIP3 and NIX for proteasomal turnover to keep levels of the receptors low, whereas TMEM11 is proposed to spatially control mitophagy by interacting with receptors at active mitophagy sites. However, it is unclear how each of these interactions is controlled and how they interplay with each other. Here, we identify a repressor of mitophagy, ARMC1, which forms a complex with TMEM11, BNIP3, and NIX. During mitophagy activation, ARMC1 dissociates from the complex, freeing the receptors to initiate mitophagy. We find that TMEM11 then acts in an antagonistic relationship with PPTC7, protecting the receptors from proteasomal degradation. Our data are consistent with a two-stage model. At steady state, a population of sentinel receptors is repressed and primed to respond to mitochondrial dysfunction. Once mitophagy is activated, TMEM11 protects BNIP3 and NIX, ensuring a sustained mitophagic response. Our findings provide a framework for understanding how two key regulatory pathways intersect to modulate receptor-mediated mitophagy.

## Introduction

Mitochondria are extraordinarily complicated organelles, with a proteome of over 1100 distinct proteins positioned within two highly organized membrane bilayers that enable the organelle to perform a vast array of crucial metabolic reactions (*1*). Because of these complex roles, mitochondria are particularly susceptible to dysfunction that ranges from the level of protein misfolding to whole-organelle defects. To deal with these issues, mitochondria have numerous quality control pathways that work to maintain organelle homeostasis (*2, 3*). One solution to dysfunction is mitophagy, whereby portions of mitochondria or entire organelles are encapsulated in autophagosomes that are targeted to the lysosome for degradation.

Several different mitophagy pathways have been identified that can be classified as ubiquitin-dependent (i.e., PINK1/Parkin) or receptor-mediated (*4*). In receptor-mediated mitophagy, outer mitochondrial membrane (OMM)-anchored proteins directly recruit autophagosome biogenesis machinery, including LC3 and WIPI, to the mitochondrial surface (*5–8*). While several mitochondrial proteins have been identified to serve as receptors, the best-characterized are BNIP3 and its paralog NIX (BNIP3L). Aside from established roles for NIX during cellular differentiation (*9–11*), both BNIP3 and NIX mediate mitochondrial quality control during stress. For example, during hypoxia, BNIP3 and NIX are transcriptionally upregulated by HIF1α to drive mitophagy (*12*). However, we and others have found that BNIP3 and NIX are also responsible for basal mitophagy in HeLa cells (*13, 14*), where Parkin is not expressed (*15*). In addition, BNIP3 and NIX have roles in tissues such as the heart and liver, where their loss is associated with the accumulation of dysfunctional mitochondria (*16, 17*).

Despite the multifunctional nature of BNIP3/NIX-mediated mitophagy, a deep understanding of their post-translational regulation is still lacking. BNIP3 and NIX are both phosphorylated at numerous sites, though the specific roles of these modifications are still unclear (*18–24*). The proteins also form SDS-resistant dimers, which is thought to be a contributor to their activation (*25*). It was recently revealed by the collective work of several labs that BNIP3 and NIX also undergo rapid turnover by the proteasome as a mechanism to keep levels of the receptors low (*12*). This pathway works through the coordinated action of PPTC7, a mitochondrial-targeted phosphatase, and FBXL4, an F-box component of the SKP-Cullin-F-box E3 ubiquitin ligase complex (*26*). While a subset of PPTC7 is imported into the mitochondrial matrix, a portion of the protein remains on the OMM where it binds to BNIP3 and NIX, forming a scaffold to mediate recruitment of FBXL4 and promote proteasomal turnover of the receptors. Loss of either PPTC7 or FBXL4 leads to marked increases in protein levels of BNIP3 and NIX, elevated levels of mitophagy, and reduced mitochondrial content (*18, 27–34*).

In addition to the recent work identifying the role of PPTC7 and FBXL4 in BNIP3/NIX turnover, we recently identified an OMM protein involved in BNIP3/NIX regulation, TMEM11 (*13*). TMEM11 forms a high molecular weight complex with BNIP3 and NIX, and during pseudohypoxic treatments that induce mitophagy, TMEM11 co-enriches with BNIP3 and NIX along with LC3 at discrete mitophagy sites on the OMM. Depletion of TMEM11 leads to increased BNIP3/NIX-dependent mitophagic flux, though the mechanism for this is unknown. TMEM11 additionally forms interactions with the Mitochondrial Intermembrane space Bridging (MIB) complex that encompasses the cristae-organizing MICOS complex on the inner membrane and OMM proteins including the β-barrel assembly SAM complex (*13, 35*). This led us to previously speculate that, via its interaction with the MIB complex, TMEM11 spatially regulates mitophagy. Despite this, the role of TMEM11 in mitophagy regulation remains unclear, and if and how TMEM11 interfaces with the PPTC7/FBXL4 turnover pathway has not been interrogated.

Here, we explore a functional role of TMEM11 in mitophagy. We focus on a recently characterized interaction between TMEM11 and the OMM Armadillo repeat containing protein ARMC1 (*36*), finding that ARMC1 is part of a TMEM11-BNIP3-NIX complex. However, our data suggest that during mitophagy induction, ARMC1 dissociates from the complex. Loss- and gain-of-function studies suggest that through its association with TMEM11, ARMC1 acts to suppress basal mitophagic flux. Further, our data indicate that once mitophagy is activated, TMEM11 protects BNIP3 and NIX from PPTC7/FBXL4-mediated proteasomal degradation. Our findings are consistent with a two-stage model of mitophagy, where low levels of a population of sentinel receptors are kept inactive prior to mitophagy induction, followed by an active stage where BNIP3 and NIX are protected from premature degradation.

## Results

### ARMC1 and TMEM11 associate in a complex with BNIP3 and NIX

Recently, TMEM11 was determined to interact with ARMC1 as part of an OMM complex whose members include mitochondrial morphology-regulating MTFR proteins (MTFR1, MTFR1L, MTFR2) and the mitochondrial motility MIRO proteins (MIRO1, MIRO2) (*36*). Via this complex, ARMC1 is proposed to control subcellular mitochondrial distribution. Because TMEM11 additionally forms a complex with BNIP3 and NIX, this raised the question of whether the ARMC1-TMEM11 association could be involved in mitophagy regulation. Consistent with this possibility, ARMC1 was one of the top identified proteins in addition to BNIP3 and NIX in our previous TMEM11 interactome analysis (*13*). To validate this finding and confirm that the association between ARMC1 and TMEM11 was generalizable, we prepared crosslinked extracts from U2OS osteosarcoma cells depleted of endogenous TMEM11 by CRISPRi and expressing APEX2-GFP-TMEM11 at close to endogenous levels (*13*). Subsequent immunoprecipitation (IP)/western analysis with GFP antibody indicated that APEX2-GFP-TMEM11 was able to robustly pull down BNIP3 and NIX, as well as ARMC1, though not the OMM translocon subunit TOMM20 (Fig. 1A).

**Figure 1.**
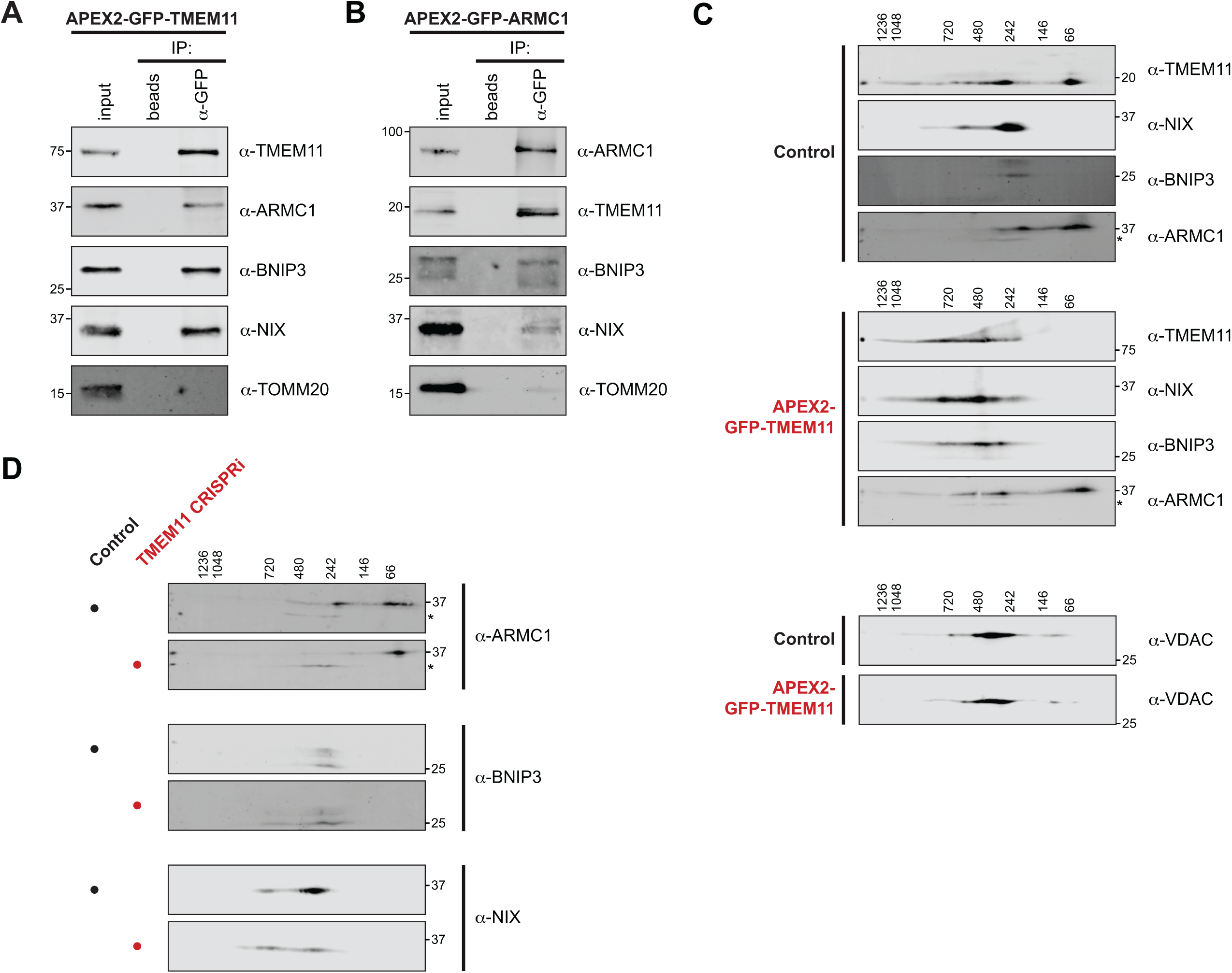
ARMC1 and TMEM11 associate in a complex with BNIP3 and NIX. **(A)** Western blots with the indicated antibodies are shown of immunoprecipitations (IPs) from cross-linked extracts from U2OS TMEM11 CRISPRi cells expressing APEX2-GFP-TMEM11. 2% of the total input and 10% of the eluate were loaded. Blots shown are representative of two independent experiments. **(B)** As in (A) for U2OS ARMC1 CRISPRi cells expressing APEX2-GFP-ARMC1. **(C)** Comparative two-dimensional Blue Native PAGE (2D-PAGE) analysis is shown with the indicated antibodies from digitonin-solubilized mitochondria isolated from U2OS CRISPRi control cells (top) or U2OS TMEM11 CRISPRi cells expressing APEX2-GFP-TMEM11 (middle). A comparison of VDAC migration pattern in each condition is shown at bottom. Asterisks indicate a cross-reacting band. Molecular weight markers displayed at top indicate native gel sizing. Blots shown are representative of two independent experiments. **(D)** As in (C) for U2OS control CRISPRi cells (indicated by black circles) or U2OS TMEM11 CRISPRi cells (red circles). See also Figures S1-S3.

To confirm the interaction between ARMC1 and TMEM11 could be detected reciprocally and test for a potential interaction with BNIP3 and NIX, we generated U2OS cells where endogenous ARMC1 was depleted using CRISPRi (Fig. S1A) and APEX2-GFP-ARMC1 was stably expressed at close to endogenous levels (Fig. S1B). APEX2-GFP-ARMC1 localized to mitochondria and was able to rescue the mitochondrial morphology defects of cells stably depleted of ARMC1 (Fig. S1C-S1D), indicating that the tagging of ARMC1 did not interfere with its function, in agreement with published observations (*36*). We then performed crosslinking IP/western analysis of APEX2-GFP-ARMC1 expressing cells with GFP antibody and identified a robust association between ARMC1 and TMEM11 (Fig. 1B). In addition, we also detected BNIP3 as well as a small subset of NIX in our IPs (Fig. 1B), suggesting that ARMC1 associates with the mitophagy receptors in addition to TMEM11.

To interrogate whether ARMC1 was part of the high molecular weight complex of TMEM11, BNIP3, and NIX that we previously observed (*13*), we performed two-dimensional Blue Native PAGE (2D BN-PAGE) analysis. We isolated a crude mitochondrial fraction from wild-type U2OS CRISPRi cells and solubilized it with the mild detergent digitonin prior to electrophoresis. Consistent with our previous results (*13*), a subset of TMEM11 protein associated in an ∼250 kD complex, similar to BNIP3 and NIX, while a portion of all three proteins was found in higher molecular weight complexes (Fig. 1C, top panel). A substantial portion of ARMC1 protein also migrated at ∼250 kD, though by comparison very little of the protein was found at higher molecular weights (Fig. 1C).

Previously, to determine if the 2D BN-PAGE distribution pattern of TMEM11, BNIP3, and NIX indicated their association in a complex, we compared their migration pattern when TMEM11 was fused to high molecular weight tags, which caused an apparent and selective shift of all three proteins to a larger size (*13*). To include ARMC1 in a similar analysis, we isolated crude mitochondria from U2OS CRISPRi cells depleted of endogenous TMEM11 and expressing APEX2-GFP-TMEM11. The fusion of high molecular weight tags (∼60 kD) to TMEM11 (endogenous size of ∼20 kD) led to a specific increase in the size of complexes of BNIP3 and NIX (Fig. 1C, middle panel). The enlarged size of TMEM11 also corresponded to an altered migration pattern of ARMC1 towards higher molecular weights (Fig. 1C, compare top and middle panels). Notably, the shifted population of ARMC1 co-migrated with BNIP3 and NIX. In contrast, and as expected (*13*), while the abundant OMM VDAC proteins migrate in similar molecular weight complexes to BNIP3 and NIX, their migration pattern was unaffected by the increased size of TMEM11 (Fig. 1C, bottom panel). These data indicate that the portion of ARMC1 that associates with TMEM11 can also reside in a complex with BNIP3 and NIX.

Using a yeast two-hybrid assay, we previously determined that TMEM11 could directly associate with BNIP3 and NIX (*13*). We therefore examined potential interactions with ARMC1 using this approach. In line with our IP/western and 2D BN-PAGE analyses, we identified direct interactions between ARMC1, BNIP3, NIX, and TMEM11 (Fig. S2). However, neither TMEM11 nor ARMC1 interacted with the known BNIP3/NIX interactor PPTC7 (Fig. S2) (*18*), highlighting the specificity of these interactions and suggesting they may regulate BNIP3/NIX as part of distinct complexes.

We next asked whether the association of ARMC1 with BNIP3 and NIX in higher molecular weight complexes occurs in a TMEM11-dependent manner. As ARMC1 peripherally associates with mitochondria through a C-terminal domain that directly associates with TMEM11 (*36*), we considered that ARMC1 may fail to associate with mitochondria in the absence of TMEM11. However, in TMEM11 CRISPRi cells, ARMC1 levels were unaffected, and by immunofluorescence microscopy, the protein retained its normal distribution pattern on the OMM (Fig. S3). We then purified mitochondria from control and TMEM11 CRISPRi cells and performed comparative 2D BN-PAGE. While TMEM11 depletion minimally affected BNIP3 and NIX migration, consistent with our previous results (*13*), the population of ARMC1 that associates with TMEM11, BNIP3, and NIX (∼250 kD) was drastically reduced in the absence of TMEM11 (Fig. 1D). In total, our data indicate that TMEM11 directly mediates an association between ARMC1 and BNIP3 and NIX in high molecular weight complexes.

### ARMC1 dissociates from TMEM11/BNIP3/NIX complexes during mitophagy induction

A common paradigm for induction of BNIP3/NIX-dependent mitophagy is hypoxia or pseudohypoxia treatments, whereby HIF1α stabilization drives BNIP3 and NIX transcription and the formation of BNIP3/NIX mitophagy sites on the OMM (*13, 34, 37*). Previously, we found that most of these BNIP3/NIX sites were co-enriched for TMEM11, indicative of a role for TMEM11 at sites of mitophagy. We therefore wondered whether ARMC1 co-distributes with TMEM11 during induced mitophagy. To visualize mitophagy sites without affecting their behavior by protein overexpression, we used CRISPR to chromosomally tag TMEM11 with mScarlet in U2OS cells (Fig. S4A). Importantly, TMEM11 function was not apparently impacted, as cells maintained normal mitochondrial morphology as compared to TMEM11-depleted cells (Fig. S4B-S4C). As expected, pseudohypoxia treatment with CoCl_2_ caused redistribution of mScarlet-TMEM11 to discrete foci on the mitochondrial membrane that co-enriched with BNIP3 relative to the OMM protein TOMM20 (Fig. S5A). A quantitative analysis found that 79% of BNIP3-enriched foci associated with the OMM exhibited a 2-fold or greater enrichment of TMEM11, similar to our previous results (Fig. S5C) (*13*). Similar results were observed with iron-chelation mediated stabilization of HIF1α using deferiprone (DFP) (88% co-enrichment; Fig. 2A, Fig. 2C), which we use going forward due to its more homogenous effect on BNIP3 levels across cells.

**Figure 2.**
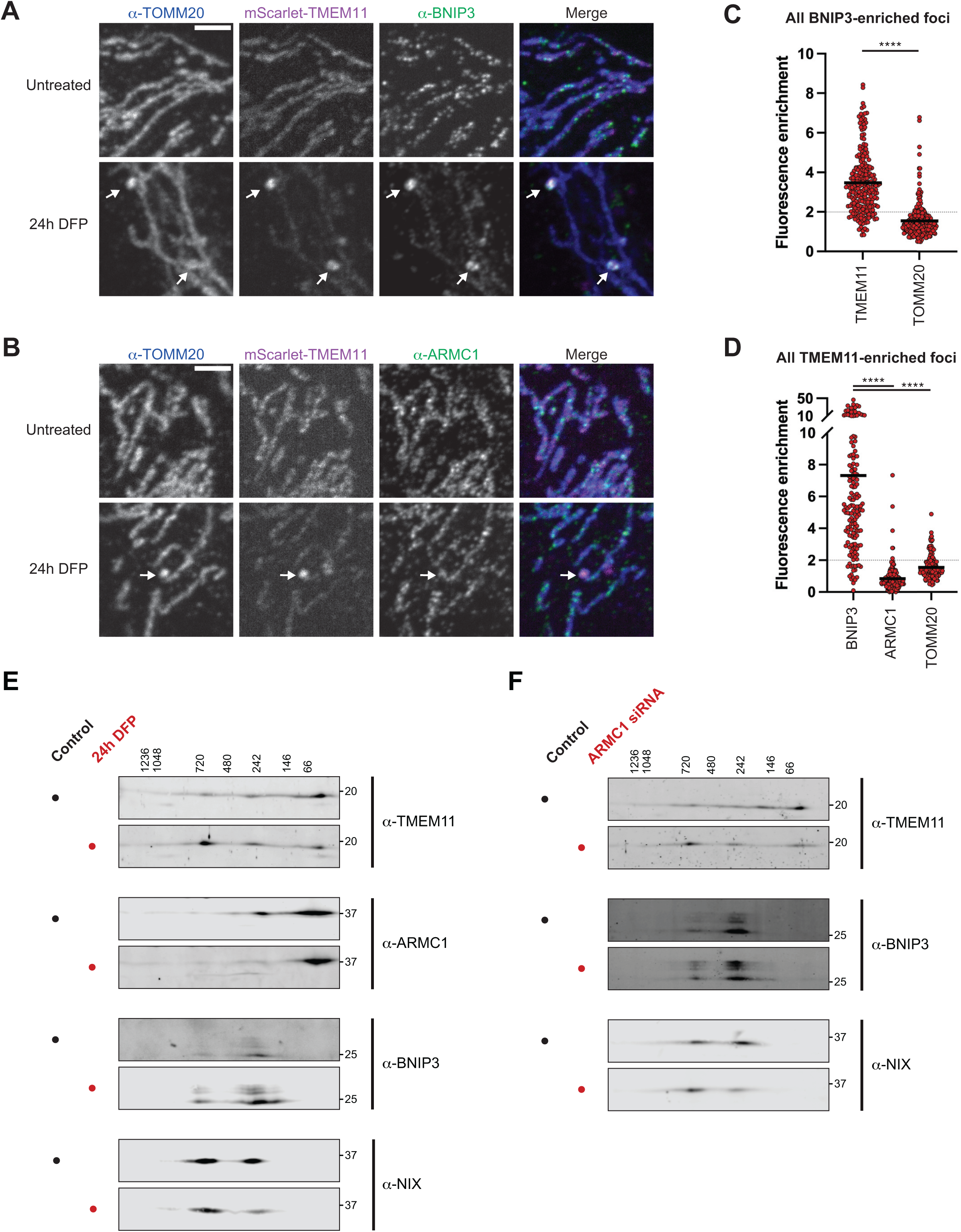
ARMC1 dissociates from TMEM11/BNIP3/NIX complexes during mitophagy induction. **(A-B)** Single plane confocal images are shown of U2OS mScarlet-TMEM11 CRISPR knock-in cells that were fixed and stained for TOMM20 and BNIP3 (A) or ARMC1 (B) before and after 1 mM DFP treatment for 24h. Arrows mark BNIP3 and/or TMEM11-enriched mitophagosomes. Scale bars = 3 µm. **(C)** A graph is shown of the relative fluorescence enrichment of TMEM11 or TOMM20 at BNIP3-enriched foci relative to signal at nearby mitochondrial tubules from images as in (A). Horizontal lines mark means for each condition. Data represent n=245 foci collected in three independent experiments. **(D)** A graph is shown as in (C) of the relative fluorescence enrichment of BNIP3, ARMC1, and TOMM20 at all mScarlet-TMEM11 enriched foci from images as in (A-B). Data represent n=153 foci (BNIP3) and n=165 foci (ARMC1, TOMM20) collected in three independent experiments. **(E)** Comparative 2D BN-PAGE analysis is shown with the indicated antibodies from digitonin-solubilized mitochondria isolated from wild-type U2OS cells before (black circles) or after (red circles) treatment with 300 µM DFP for 24h. Molecular weight markers displayed at top indicate native gel sizing. Blots shown are representative of three independent experiments. **(F)** As in (E) for mitochondria isolated from U2OS cells transfected with scrambled control siRNA (black circles) or ARMC1 siRNA #1 (red circles). Blots shown are representative of two independent experiments. Asterisks (****p<0.0001) represent a paired t-test (C) or one-way ANOVA with Tukey’s multiple comparisons test (D). See also Figures S4-S7.

With mScarlet-TMEM11 cells in hand, we treated them with DFP to induce mitophagy, and then fixed and stained cells with antibodies against endogenous ARMC1 and TOMM20. In contrast to the strong co-enrichment of BNIP3 at TMEM11-enriched sites induced by DFP (83% BNIP3-enriched), ARMC1 was rarely co-enriched with TMEM11 (4% of foci) and was frequently de-enriched relative to TOMM20 (Fig. 2B and Fig. 2D). Unlike BNIP3 and NIX, neither TMEM11 nor ARMC1 levels drastically changed after DFP treatment, indicating the difference between ARMC1 and TMEM11 distribution is not due to changes in their relative abundance (Fig. S6). Our findings were generalizable to other paradigms of pseudohypoxia, as ARMC1 also did not co-enrich with TMEM11-marked enrichments after CoCl_2_ treatment (Fig. S5B, S5D). Thus, even though ARMC1 associates with a TMEM11/BNIP3/NIX complex, the protein does not co-enrich with TMEM11 at mitophagy sites.

We next determined whether the distribution of proteins amongst complexes changed during the induction of mitophagy. We isolated mitochondria from U2OS cells before and after treatment for 24h with DFP and performed comparative 2D BN-PAGE analysis of TMEM11, ARMC1, BNIP3, and NIX. The distribution pattern for ARMC1 shifted, with very little of the protein remaining at ∼250 kD after DFP treatment (Fig. 2E). Instead, ARMC1 was found primarily at a lower molecular weight, similar to the distribution pattern observed in TMEM11-depleted cells (compare Fig. 2E to Fig. 1D). In addition, we observed changes in the amount of TMEM11 and NIX in each complex size, with higher levels of each protein in a ∼700 kD complex relative to the ∼250 kD complex after DFP treatment (Fig. 2E). These data, in combination with our confocal microscopy analysis, suggest that ARMC1 dissociates from the TMEM11/BNIP3/NIX complex during mitophagy induction. In addition, the relative increase in ∼700 kD complexes after DFP treatment suggests that these may contain a more active form of a mitophagy complex.

Given that ARMC1 appeared to dissociate from the complex of TMEM11, BNIP3, and NIX upon DFP treatment, we asked how ARMC1 depletion impacted the distribution of each of the other proteins across complexes. Mitochondria were isolated from U2OS cells depleted of ARMC1 by siRNA (Fig. S7A) and 2D BN-PAGE analysis was performed in comparison to cells treated with scrambled control siRNA (Fig. 2F). ARMC1 depletion led to subtle increases in BNIP3 and NIX levels (Fig. S7A) and both TMEM11 and NIX were found more frequently in a ∼700 kD complex after ARMC1 depletion, similar to our observations in DFP-treated mitochondria (compare Fig. 2E and 2F). These data indicate that ARMC1 is not required for the formation of TMEM11/BNIP3/NIX complexes and that depletion of ARMC1 phenocopies aspects of mitophagy induction.

### ARMC1 levels inversely correlate with basal mitophagic flux

Because ARMC1 dissociates from TMEM11/BNIP3/NIX during pseudohypoxic mitophagy induction, we hypothesized that ARMC1 plays an inhibitory role in mitophagy. Previously, we observed that in HeLa cells, low levels of BNIP3/NIX-dependent basal mitophagic flux could be observed using the reporter mito-mKeima (*13*). mKeima is a fluorophore that exhibits a spectral shift in acidic environments, and when targeted to mitochondria, can be used to monitor the presence of the organelle in the acidic lysosome (*38*). To address if ARMC1 levels affect basal mitophagic flux, we transfected HeLa mito-mKeima cells with either a scrambled control or two independent siRNAs targeting ARMC1 (Fig. S7B) and imaged by confocal microscopy. We then assessed basal mitophagy by counting the number of acidified mitochondrial puncta per cell blind to sample identity. In cells depleted of ARMC1, basal mitophagy was significantly elevated compared to control cells (Fig. 3A-3B). We evaluated the levels of BNIP3 and NIX after siRNA treatment, and unlike in U2OS cells, the total levels of the receptors were not substantially altered (Fig. S7B). Co-transfection with siRNAs targeting BNIP3 and NIX reduced levels of basal mitophagy, consistent with previous findings (*13, 14*) and indicating that the mitophagy increase in the absence of ARMC1 was dependent on the BNIP3/NIX pathway (Fig. 3A-3B, S7B).

**Figure 3.**
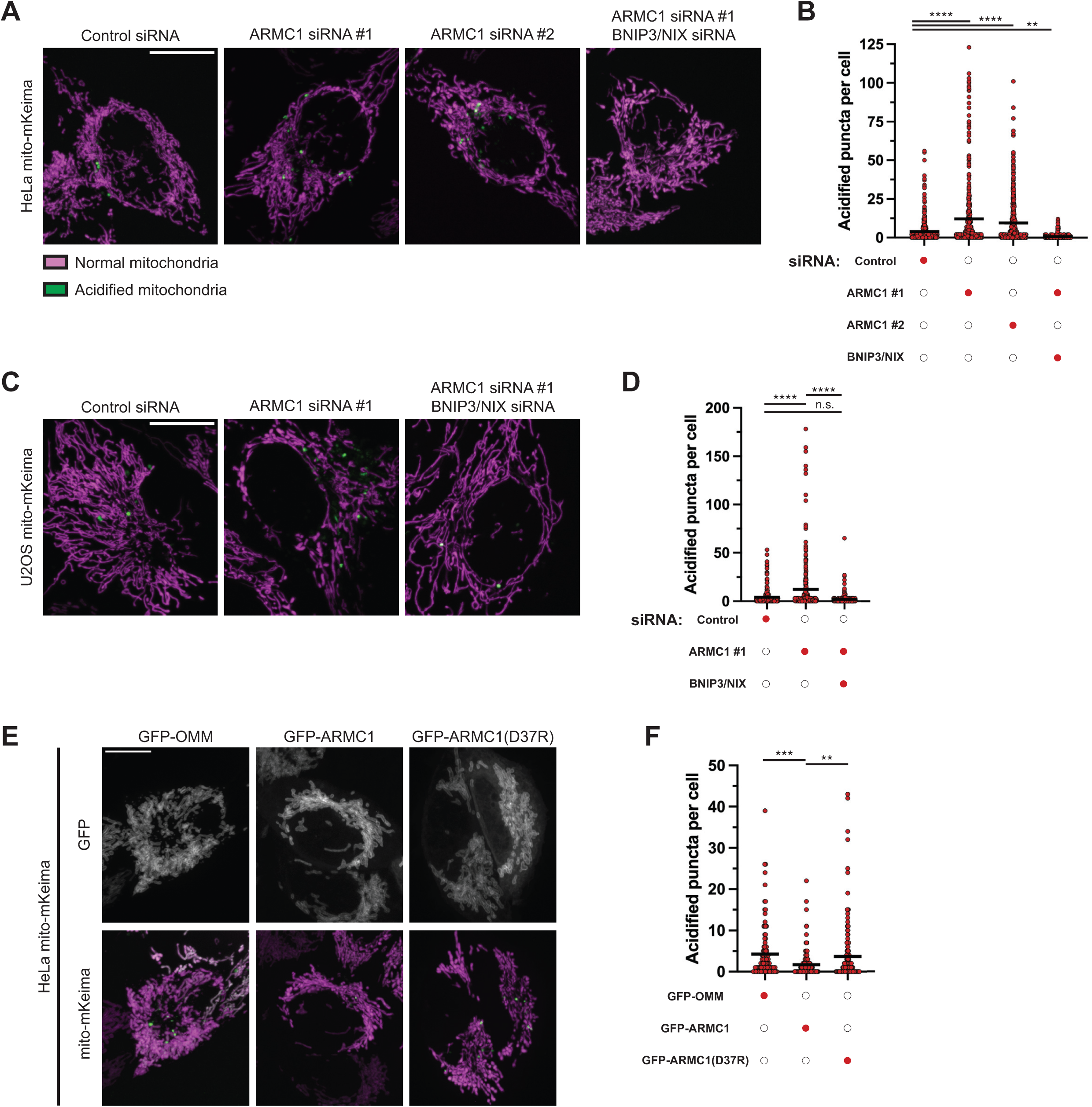
ARMC1 levels inversely correlate with basal mitophagic flux. **(A, C)** Merged maximum intensity projection confocal images are shown of representative HeLa mito-mKeima (A) or U2OS mito-mKeima (C) expressing cells treated with the indicated siRNAs. Images are pseudocolored to represent mitochondria with a neutral pH (magenta; 488 nm laser excitation) and acidified mitochondrial signal (green; 561 nm laser excitation). **(B, D)** Graphs are shown of quantifications of the number of acidified mitochondria puncta per cell in the indicated siRNA-treated conditions from cells as in (A, C). Data represent approximately 300-400 cells per condition collected in four independent experiments. Horizontal lines mark means for each condition. **(E)** Representative maximum intensity projection confocal images are shown from HeLa cells stably expressing mito-mKeima and transiently transfected with the indicated GFP fusions. GFP signal is shown at top and mito-mKeima signal displayed as in (A) is shown at bottom. **(F)** A graph is shown as in (B, D) of cells transfected with comparable levels of the indicated GFP fusion constructs as in (E). Data represent approximately 100-200 cells per condition collected in three independent experiments. Asterisks (****p<0.0001, ***p<0.001, **p<0.01) represent one-way ANOVA with Tukey’s multiple comparisons test. N.S. indicates not statistically significant. Scale bars (A, C) = 15 µm; (E) = 10 µm. See also Figure S7.

To next determine if this finding extended beyond HeLa cells, we treated U2OS mito-mKeima cells with siRNA targeting ARMC1. Similar to our findings in HeLa, U2OS mitophagic flux increased in the absence of ARMC1 (Fig. 3C-3D; Fig. S7C). The increase in mitophagy levels was completely diminished by combined depletion of ARMC1, BNIP3, and NIX, indicating that ARMC1 depletion increases mitophagic flux via the BNIP3/NIX pathway (Fig. 3C-3D; Fig. S7C). These results are consistent with a potential role for ARMC1 as a repressor of mitophagy.

Because basal mitophagic flux in HeLa cells is BNIP3 and NIX-dependent and ARMC1 depletion is sufficient to increase basal mitophagy, we asked if an increase in ARMC1 levels is sufficient to reduce basal mitophagy. To address this, we transiently transfected HeLa mito-mKeima cells with GFP-ARMC1, or, as a control, GFP fused to the transmembrane domain of the tail-anchored OMM protein FIS1 (hereafter GFP-OMM). In cells with comparable levels of GFP expression, basal mitophagy was significantly lessened by overexpression of GFP-ARMC1 (Fig. 3E-3F). This decrease was not due to a reduction in BNIP3 or NIX levels, which were unaffected by overexpression of ARMC1 (Fig. S7D). To determine if the decrease in basal mitophagy was dependent on the association of ARMC1 with TMEM11, we compared mitophagic flux in GFP-ARMC1 transfected cells to that of a mutant of GFP-ARMC1, D37R, that is unable to bind to TMEM11 (*36*). However, unlike wild-type ARMC1, the D37R mutant had no effect on mitophagy (Fig. 3E-3F). These data indicate that an increase in ARMC1 levels is sufficient to repress basal mitophagy in a manner dependent on the ability of ARMC1 to bind to TMEM11.

### Overexpression of TMEM11 increases BNIP3/NIX levels and drives basal mitophagy

Loss of ARMC1 led to an increase in basal mitophagic flux, while its overexpression decreased mitophagy, consistent with a potential repressive role. Because TMEM11 localizes to mitophagy sites independently of ARMC1, we predicted that elevated TMEM11 expression could bypass ARMC1-mediated repression and increase basal mitophagic flux. Notably, in HeLa cells, ARMC1 is estimated to have similar protein levels to TMEM11 (*36, 39*). We transiently transfected HeLa mito-mKeima cells with GFP-OMM or GFP-TMEM11 and quantitatively assessed basal acidic mitochondrial puncta. Compared to either an empty vector control or GFP-OMM, overexpression of GFP-TMEM11 significantly increased basal mitophagic flux (Fig. 4A-4B).

**Figure 4.**
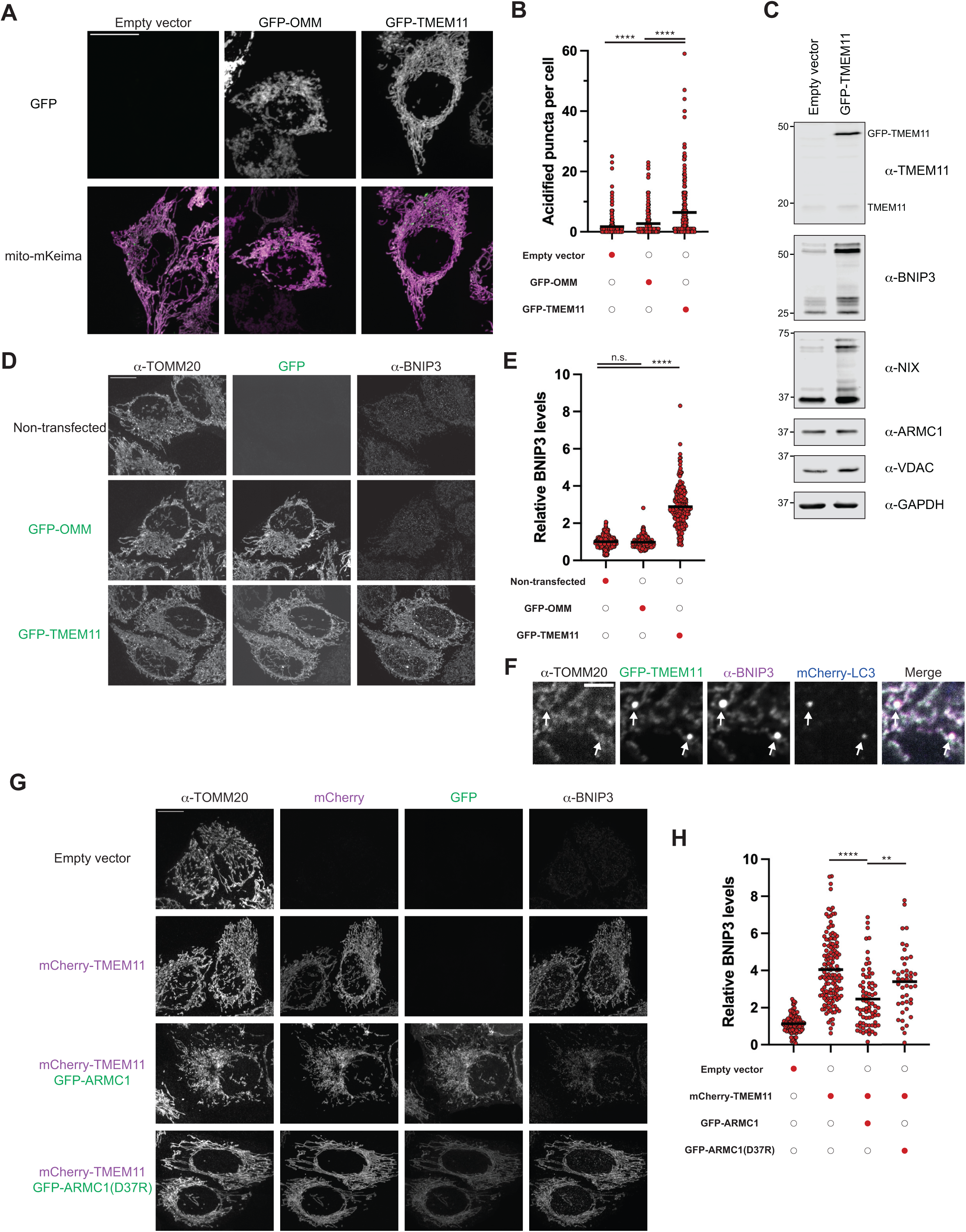
Overexpression of TMEM11 increases BNIP3/NIX levels and drives basal mitophagy. **(A)** Representative maximum intensity projection confocal images are shown from HeLa cells stably expressing mito-mKeima and transiently transfected with the indicated GFP fusions. GFP signal is shown at top and mito-mKeima signal (pseudo-colored to indicate mitochondrial with a neutral pH (magenta) and acidic pH (green)) is shown at bottom. **(B)** A graph is shown of the number of acidified mitochondria puncta per cell from cells transfected with empty vector or comparable levels of the indicated GFP fusion constructs as in (A). Horizontal lines mark means for each condition. Data represent approximately 200-300 cells per condition collected in three independent experiments. **(C)** Western blots are shown with the indicated antibodies of lysate from HeLa cells transiently transfected with an empty vector or GFP-TMEM11. **(D)** Representative maximum intensity projection confocal images are shown of HeLa cells transiently transfected with GFP-OMM or GFP-TMEM11 and fixed and stained for BNIP3 and TOMM20. A non-transfected cell is shown at top for comparison. Unmerged images are shown to indicate the signal in each channel, and images of GFP and BNIP3 were identically contrasted for all conditions. **(E)** A graph showing a quantification of the relative BNIP3 levels in individual cells, as in (D), within a consistent range of GFP expression for each condition. The mean BNIP3 signal was determined for all non-transfected cells in each experimental replicate and all individual cell values are shown relative to the respective mean. Data represent approximately 200-300 cells per condition collected in three independent experiments. **(F)** A single plane confocal image is shown of a HeLa cell transiently transfected with GFP-TMEM11 and mCherry-LC3 and fixed and stained with BNIP3 and TOMM20. Arrows mark sites of LC3-marked TMEM11/BNIP3-enriched mitophagosomes. **(G)** Representative maximum intensity projection confocal images are shown of HeLa cells transiently transfected with the indicated constructs and fixed and stained for BNIP3 and TOMM20. Images of GFP, mCherry, and BNIP3 were identically contrasted for all conditions. **(H)** As in (E) for cells imaged as in (G). Cells were analyzed within a consistent range of GFP and/or mCherry expression between conditions and data represent at least n=44 cells per condition collected in three independent experiments. Relative BNIP3 levels were determined relative to the mean of the empty vector control for each experimental replicate. Asterisks (****p<0.0001, **p<0.01) represent one-way ANOVA with Tukey’s multiple comparisons test. N.S. indicates not statistically significant. Scale bars: (A, D, G) = 15µm; (F) 3µm.

We considered whether the increase in mitophagy in cells overexpressing TMEM11 could be explained by alterations in the levels of ARMC1, BNIP3, or NIX. While ARMC1 levels were unaffected, BNIP3 and NIX levels were substantially higher in HeLa cells transiently transfected with GFP-TMEM11 (Fig. 4C). To assess BNIP3 levels specifically in GFP-TMEM11 overexpressing cells, we performed immunofluorescence with BNIP3 and TOMM20 antibodies and imaged by confocal microscopy. Consistent with western analysis, BNIP3 levels were significantly higher specifically in cells overexpressing TMEM11 in comparison to GFP-OMM or non-transfected cells (Fig. 4D-4E). We also routinely observed TMEM11 and BNIP3 co-enrichment at discrete foci in GFP-TMEM11 overexpressing cells that co-localized with mCherry-LC3 (Fig. 4F), consistent with the increased basal mitophagic flux in these cells. To determine if GFP-TMEM11 overexpression also led to an increase in BNIP3 and NIX levels in U2OS cells, we utilized CRISPRi cells with endogenous TMEM11 depleted and transduced with a GFP-TMEM11 expressing plasmid (*13*). As in HeLa cells, GFP-TMEM11 overexpression in U2OS cells increased the levels of BNP3 and NIX without affecting the levels of ARMC1 (Fig. S8A).

Our data are consistent with the possibility that TMEM11 overexpression increases BNIP3 and NIX levels and mitophagy due to altered stoichiometry between ARMC1 and TMEM11. We thus asked whether overexpression of ARMC1 could neutralize the effect of TMEM11 overexpression on the levels of BNIP3 and NIX. We transiently co-transfected HeLa cells with mCherry-TMEM11 and GFP-ARMC1 or GFP-ARMC1(D37R), and fixed and immunolabeled cells for BNIP3 and TOMM20. We then quantitatively assessed relative BNIP3 levels in cells expressing similar levels of each construct. As with GFP-tagged TMEM11, mCherry-TMEM11 expression significantly increased the levels of BNIP3 (Fig. 4G-4H). However, co-expression with GFP-ARMC1 diminished the effect of TMEM11 overexpression in comparison to the ARMC1 D37R variant (Fig. 4G-4H). Together, these data suggest that TMEM11 overexpression promotes an increase in BNIP3 levels by outcompeting its association with ARMC1.

### TMEM11 promotes stabilization of BNIP3 and NIX

The increase in BNIP3 and NIX levels in cells overexpressing TMEM11 phenocopies cells depleted of FBXL4 and PPTC7, factors responsible for proteasomal turnover of BNIP3 and NIX (*12*). We thus hypothesized that TMEM11 impedes the turnover of BNIP3 and NIX and asked if increased TMEM11 levels promoted BNIP3 and NIX protein stability. We pre-treated U2OS CRISPRi control cells or those overexpressing GFP-TMEM11 (Fig. S8A) with DFP for 24h to increase the levels of BNIP3 and NIX to more easily monitor protein turnover. We then monitored protein stability during a 5-hour treatment of cycloheximide to block new protein synthesis. In control cells, BNIP3 and NIX each exhibited a half-life of less than two hours (Fig. 5A-5B). In comparison, only minimal turnover of the receptors was observed within 5 hours of cycloheximide treatment in cells overexpressing TMEM11 (Fig. 5A-5B). As an orthogonal approach, we monitored the accumulation of BNIP3 and NIX in control and TMEM11-overexpressing cells upon treatment with the proteasomal inhibitor MG-132. In agreement with previous observations (*34*), MG-132 treatment in otherwise untreated cells led to the accumulation of BNIP3 and NIX, suggesting they are basally turned over by the proteasome. While TMEM11 overexpression led to a substantial increase in BNIP3 and NIX levels, the proteins did not further accumulate after treatment with MG-132 (Fig. S9A-S9B). Together, these data suggest that elevated TMEM11 levels protect BNIP3 and NIX from proteasomal degradation.

**Figure 5.**
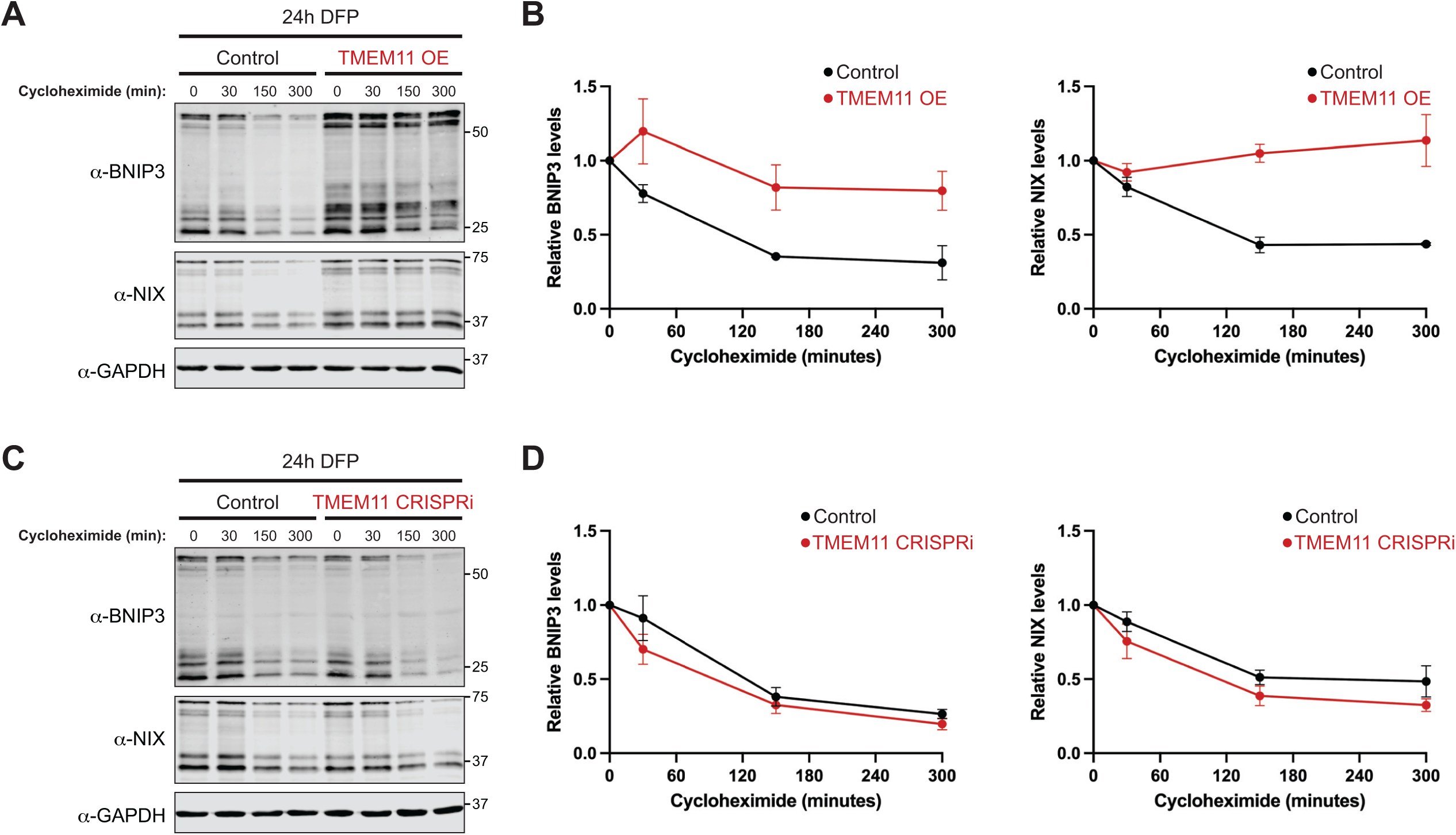
TMEM11 promotes stabilization of BNIP3 and NIX. **(A)** Representative western blots are shown with the indicated antibodies of lysates from U2OS Control CRISPRi cells (left) or U2OS TMEM11 CRISPRi cells stably expressing high levels of GFP-TMEM11 (right). Cells were treated with 1 mM DFP for 24h, followed by subsequent treatment with 50 µg/ml cycloheximide for the indicated times. **(B)** Graphs are shown of quantifications of the relative levels of BNIP3 (left) or NIX (right) after treatment with cycloheximide as in (A). The BNIP3 levels for control (black lines) and GFP-TMEM11 overexpressing cells (red lines) were normalized to GAPDH signal and determined relative to the 0-minute timepoint for each condition. Data shown are from three independent experiments and bars indicate S.E.M.. **(C-D)** As in (A-B) for U2OS CRISPRi control cells (black lines) and U2OS TMEM11 CRISPRi cells (red lines). See also Figures S8-S9.

We next asked whether the turnover of BNIP3 and NIX was accelerated in cells depleted of TMEM11. U2OS CRISPRi cells expressing control or TMEM11 sgRNAs were pre-treated with DFP for 24h prior to a time-course of cycloheximide. In comparison with control cells, the rate of turnover of BNIP3 and NIX was modestly accelerated in the absence of TMEM11 (Fig. 5C-5D). Consistent with this finding, both BNIP3 and NIX accumulated somewhat more rapidly in cells depleted of TMEM11 after MG-132 treatment (Fig S9C-S9D). Together, these data indicate that both at steady-state and after DFP treatment, TMEM11 has a protective effect against the turnover of BNIP3 and NIX.

### TMEM11 protects BNIP3 and NIX from PPTC7/FBLX4-mediated proteasomal degradation

By altering TMEM11 levels, BNIP3 and NIX are differentially susceptible to proteasomal degradation. These results led us to hypothesize that TMEM11 influences the efficacy of PPTC7 and FBXL4-mediated turnover of BNIP3 and NIX. Previously, we and others found that overexpression of PPTC7 was sufficient to reduce the levels of BNIP3 and NIX during iron-chelation-mediated pseudohypoxia (*28, 29, 34*). Because TMEM11 overexpression leads to the stabilization of BNIP3 and NIX, we wanted to determine if these factors were reciprocally antagonistic. To address this, we transiently transfected HeLa cells with PPTC7-GFP, mCherry-TMEM11, or a combination of both plasmids, followed by a 24h treatment with DFP to increase BNIP3 levels. Cells were then fixed and stained with BNIP3 and TOMM20 antibodies and imaged by confocal microscopy. As expected, overexpression of PPTC7-GFP led to a substantial loss of detectable BNIP3 staining after DFP-treatment (Fig. 6A-6B). Conversely, mCherry-TMEM11 overexpression elevated BNIP3 levels above those caused by DFP treatment, consistent with our western analysis of U2OS cells (Fig. 5A and Fig. 6B). We then examined cells transfected with both PPTC7-GFP and mCherry-TMEM11, ensuring that analyzed cells expressed similar fluorescence levels of each construct compared to singly transfected cells. Strikingly, cells overexpressing both PPTC7 and TMEM11 had similar BNIP3 levels to an empty vector control (Fig. 6A-6B). These data indicate that TMEM11 antagonizes PPTC7/FBXL4-mediated effects on BNIP3 levels.

**Figure 6.**
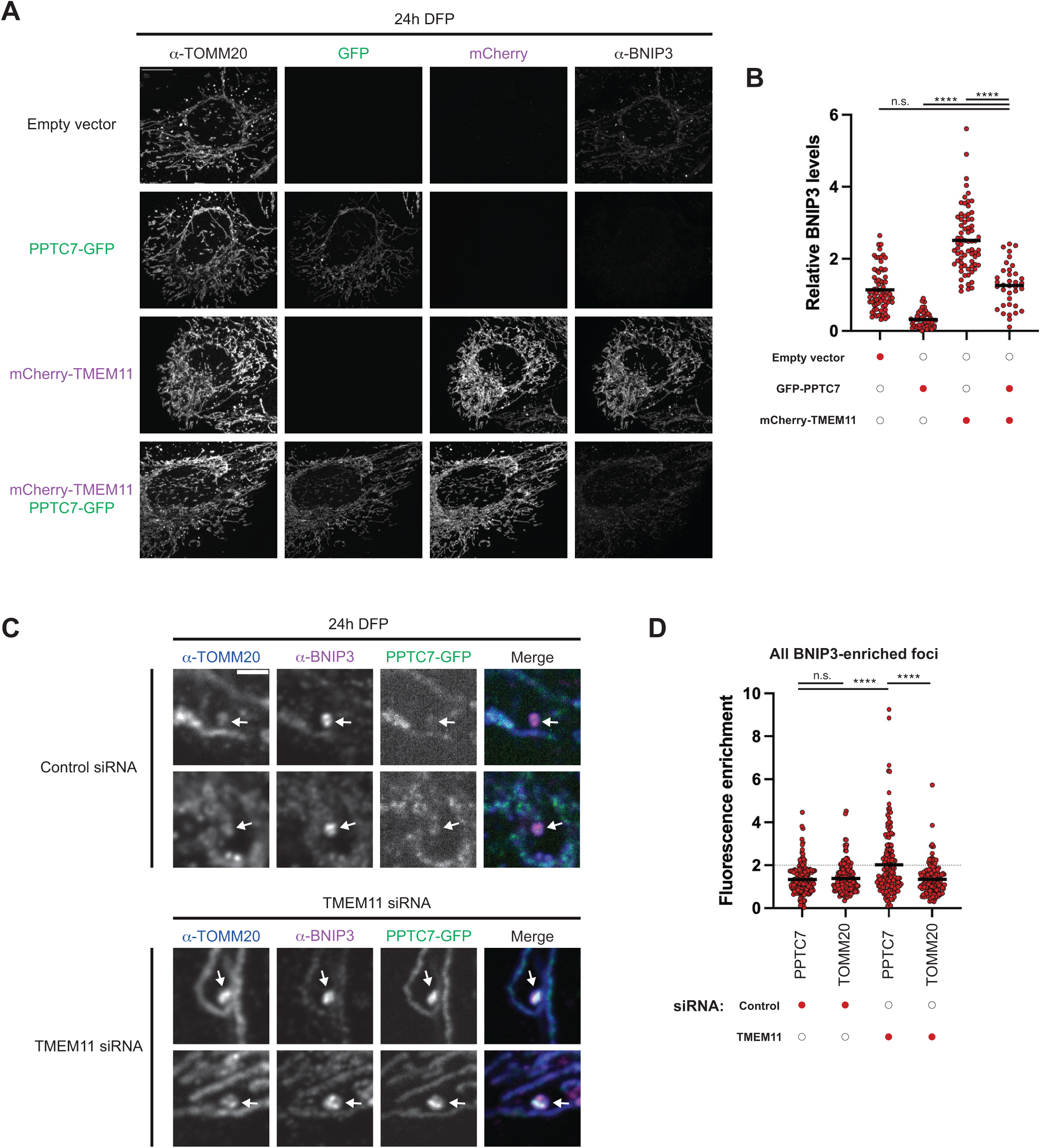
TMEM11 protects BNIP3 and NIX from PPTC7/FBLX4-mediated proteasomal degradation. **(A)** Representative maximum intensity projection confocal images are shown of HeLa cells transiently transfected with the indicated constructs, treated with 1mM DFP for 24h, and fixed and stained for BNIP3 and TOMM20. Images of GFP, mCherry, and BNIP3 were identically contrasted for all conditions. **(B)** A graph is shown of quantification of the relative BNIP3 levels in individual cells as in (A). Cells were analyzed within a consistent range of GFP and/or mCherry expression between conditions and data represent at least n=36 cells per condition collected in two independent experiments. The mean BNIP3 signal was determined for all non-transfected cells in each experimental replicate and all individual cell values are shown relative to the respective mean. **(C)** Single plane confocal images are shown of HeLa cells co-transfected with scrambled control siRNA (top) or TMEM11 siRNA (bottom) and PPTC7-GFP, treated with 1mM DFP for 24h, and fixed and stained for BNIP3 and TOMM20. Arrows mark sites of BNIP3-enriched mitophagosomes. **(D)** A graph is shown of the relative fluorescence enrichment of PPTC7 or TOMM20 at BNIP3-enriched foci relative to signal at nearby mitochondrial tubules from images as in (C). Data represent approximately n=150-200 foci per condition collected in two independent experiments. Asterisks (****p<0.0001) represent one-way ANOVA with Tukey’s multiple comparisons test. N.S. indicates not statistically significant. Scale bars: (A) = 15µm; (C) 3µm. See also Figure S8.

Our data indicate that TMEM11, either when overexpressed in excess of ARMC1, or after DFP treatment where it is dissociated from ARMC1, can serve in a protective role against PPTC7/FBXL4-mediated proteasomal turnover of BNIP3 and NIX. Previously, we examined the localization of PPTC7-GFP to BNIP3-enriched mitophagosomes induced by DFP in U2OS cells, finding they were only rarely enriched (*34*). Given that our data suggest that TMEM11 serves in a protective role against PPTC7/FBXL4-mediated BNIP3 degradation, we predicted that in the absence of TMEM11, PPTC7 would have greater accessibility to mitophagy sites. To address this, we depleted TMEM11 from U2OS cells using RNAi (Fig. S8B) and transiently expressed PPTC7-GFP at low but detectable levels so as not to promote degradation of BNIP3. We then treated cells with DFP for 24h followed by fixation and staining for BNIP3 and TOMM20 and quantitatively assessed PPTC7 enrichment at BNIP3-marked mitophagy sites. While PPTC7-GFP could be found at low levels at most mitophagy sites, it was only rarely co-enriched with BNIP3, similar to TOMM20 and consistent with our previous results (12% of sites had 2-fold enrichment; Fig. 6C-6D). Strikingly, in cells depleted of TMEM11, PPTC7 was significantly more enriched at mitophagy sites, with >2-fold enrichment at 39% of sites (Fig. 6C-6D). Because PPTC7 is dual-localized to the matrix and OMM, we considered that alterations in TMEM11 levels could impact OMM targeting of PPTC7. However, neither TMEM11 depletion nor overexpression impacted the relative levels of either isoform of PPTC7 (Fig. S8). Together, these data support the model that during mitophagy induction, TMEM11 locally protects BNIP3 at mitophagy sites from degradation mediated by PPTC7 and FBXL4.

## Discussion

Our findings suggest that ARMC1 is a novel negative regulator of BNIP3 and NIX, acting as a repressor of basal mitophagic flux through its interaction with TMEM11. During mitophagy induction, for example upon pseudo-hypoxic insults, ARMC1 dissociates from TMEM11/BNIP3/NIX, which subsequently enrich at productive mitophagy sites. There, TMEM11 plays a positive role in maintaining BNIP3/NIX protein stability, protecting the receptors from proteasomal degradation mediated by PPTC7 and FBXL4. While our previous work demonstrated increased mitophagic flux in the absence of TMEM11 (*13*), these data are likely explained by the constitutive loss of BNIP3/NIX repression by ARMC1. We find here that in the absence of TMEM11, BNIP3-marked mitophagy sites are substantially more enriched for PPTC7, indicative of reduced protection of the receptor from turnover.

Our results are consistent with a two-stage model of mitophagy regulation (Fig. 7). At steady-state basal conditions, a population of BNIP3 and NIX is targeted to the OMM and held in check by a complex of ARMC1 and TMEM11. Meanwhile, free BNIP3 and NIX are unprotected and accessible to PPTC7 and FBXL4 to mediate the rapid turnover of unbound receptors. However, once a mitochondrial insult is detected, or in the case of pseudohypoxia where mitophagy is actively stimulated, TMEM11 co-distributes with BNIP3/NIX to sites of mitophagy. There, TMEM11 ensures that BNIP3 and NIX can promote downstream mitophagy factor recruitment without interference from the proteasomal turnover pathway.

**Figure 7.**
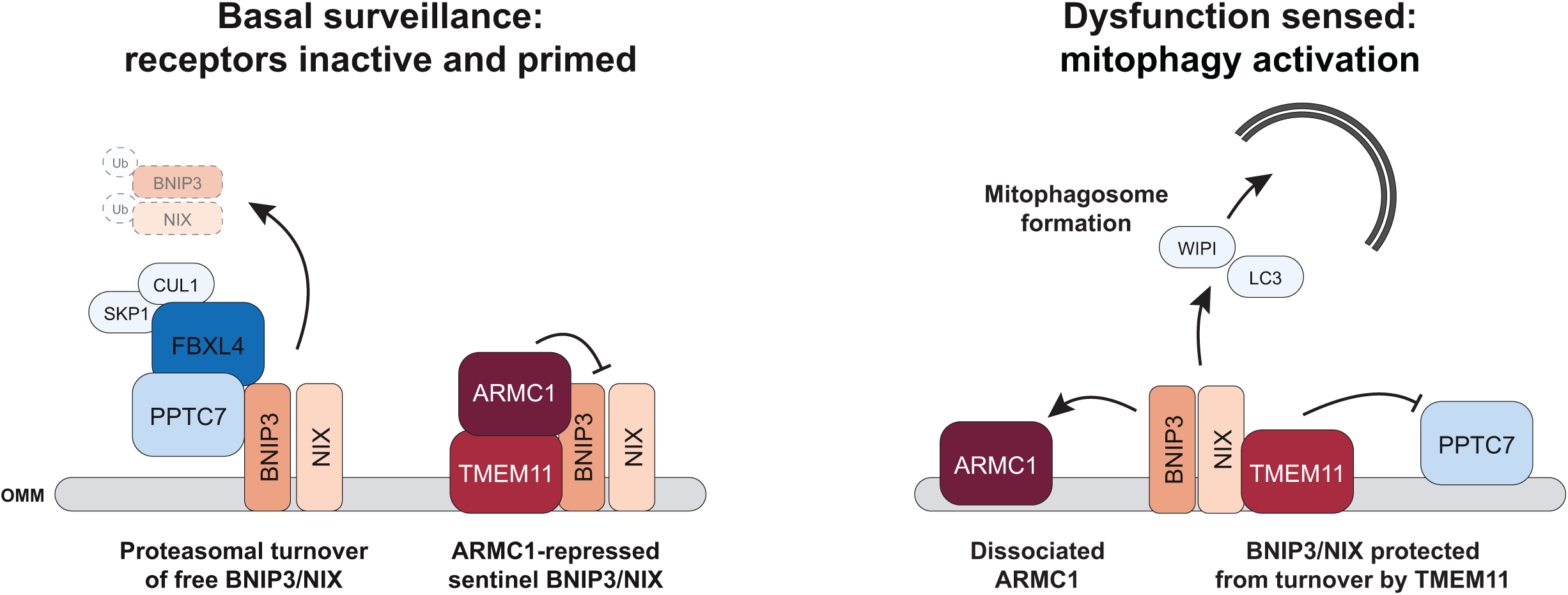
A model for the roles of ARMC1 and TMEM11 in the regulation of mitophagy. A working model is shown for the interplay between ARMC1/TMEM11 and PPTC7/FBXL4 in the regulation of BNIP3/NIX-mediated mitophagy. Under basal conditions, a subset of BNIP3 and NIX is bound by ARMC1 and TMEM11. These receptors are kept in a repressed state and can survey the outer mitochondrial membrane for mitochondrial dysfunction. BNIP3 and NIX which are not bound by ARMC1 and TMEM11 are more accessible to PPTC7, which mediates scaffolding of the FBXL4-containing SCF complex that promotes proteasomal turnover of BNIP3 and NIX. During activation of mitophagy, such as occurs during hypoxia, ARMC1 dissociates from TMEM11, freeing BNIP3 and NIX to mediate LC3 and WIPI recruitment and the formation of mitophagosomes. TMEM11 co-concentrates with BNIP3/NIX at these mitophagy sites, protecting the receptors from premature PPTC7/FBXL4-mediated turnover.

The current prevailing model for BNIP3/NIX regulation is that by continuously turning over the receptors, PPTC7 and FBXL4 prevent their activation. However, such a system necessitates time post-insult to pause turnover and allow the receptors to accumulate to sufficient levels to drive mitophagy. Additionally, the general accumulation of receptors does not enable the tight spatial control that would be necessary to sense and respond to localized mitochondrial dysfunction as would be expected of a quality control mitophagy pathway. An advantage of having a population of repressed BNIP3 and NIX is that cells can be primed and ready to sense and respond to acute mitochondrial dysfunction. Thus, we propose that ARMC1/TMEM11-bound BNIP3 and NIX serve as sentinel receptors.

A key question that arises from this model is what stress(es) mediate BNIP3 and NIX activity under basal conditions and how ARMC1 can sense and trigger the release of active BNIP3 and NIX under the protection of TMEM11. Clues into this may come from the additional interactions shared between TMEM11 and ARMC1. ARMC1 participates in two protein complexes, one with TMEM11 and a second containing DNAJC11, a member of the IMM-OMM spanning Mitochondrial Intermembrane space Bridging (MIB) complex (*36*). Likewise, TMEM11 strongly associates with members of the MIB complex, including DNAJC11 as well as the IMM cristae-organizing MICOS complex (*13, 35*). Our previous observations indicated that cristae disruption through loss of the MICOS subunit MIC60 was sufficient to drive basal BNIP3/NIX-dependent mitophagy in HeLa cells (*13*). Thus, we speculate that at least one form of dysfunction sensed through ARMC1 and/or TMEM11 is a local disruption of the MIB complex. An additional important tool in revealing the answer may lie in the distinct nature of different cell culture models. In HeLa cells where Parkin is not expressed (*15*), most basal mitophagy is dependent on BNIP3 despite its extremely low protein levels (*13, 14*). Importantly, while U2OS cells are competent for BNIP3/NIX-dependent basal mitophagy (i.e., ARMC1 depletion elicits a BNIP3/NIX-dependent response), BNIP3/NIX depletion does not diminish basal mitophagic flux. These differences may enable the identification of additional stress regimes that either universally stimulate BNIP3 or NIX, or work in specific cell types, and reveal how other mitophagy pathways may redundantly compensate for the loss of one another.

While our work provides a framework for our understanding of the functional roles of various players in mitophagy regulation, it leaves open the question of the mechanistic relationship between the key and currently identified factors. Work from Shao and colleagues identified a direct physical interaction between the ARMC1 C-terminal domain, which mediates its OMM association, and TMEM11 (*36*). AlphaFold3 modeling predicts this interaction occurs in proximity to the TMEM11 transmembrane domains (*40*). Our prior work also indicates that TMEM11 can associate with BNIP3 via their respective transmembrane domains or proximal regions (*13*). Thus, it is conceivable that a TMEM11/ARMC1/BNIP3/NIX complex, as well as the remodeling that occurs during pseudohypoxia, may be mediated at or near the membrane. An additional question raised is how the presence of TMEM11 promotes the stabilization of BNIP3 and NIX and reduces or prevents association with PPTC7. Notably, the peri-transmembrane association between TMEM11 and BNIP3 is likely distinct from that of PPTC7, which associates with BNIP3 and NIX via the cytosol-exposed BH3 and SRPE motifs of these mitophagy receptors (*28*). However, it is possible that the cytosolic domain of TMEM11 sterically hinders PPTC7 association. Future work to precisely define these mechanisms will be necessary to gain a more complete understanding of mitophagy regulation. However, our work provides important insights into the dynamic interplay between mitophagy receptor repression, activation, and turnover, which we believe forms the basis of understanding how BNIP3/NIX mitophagy is utilized to maintain mitochondrial homeostasis.

---

**Figure S1.**
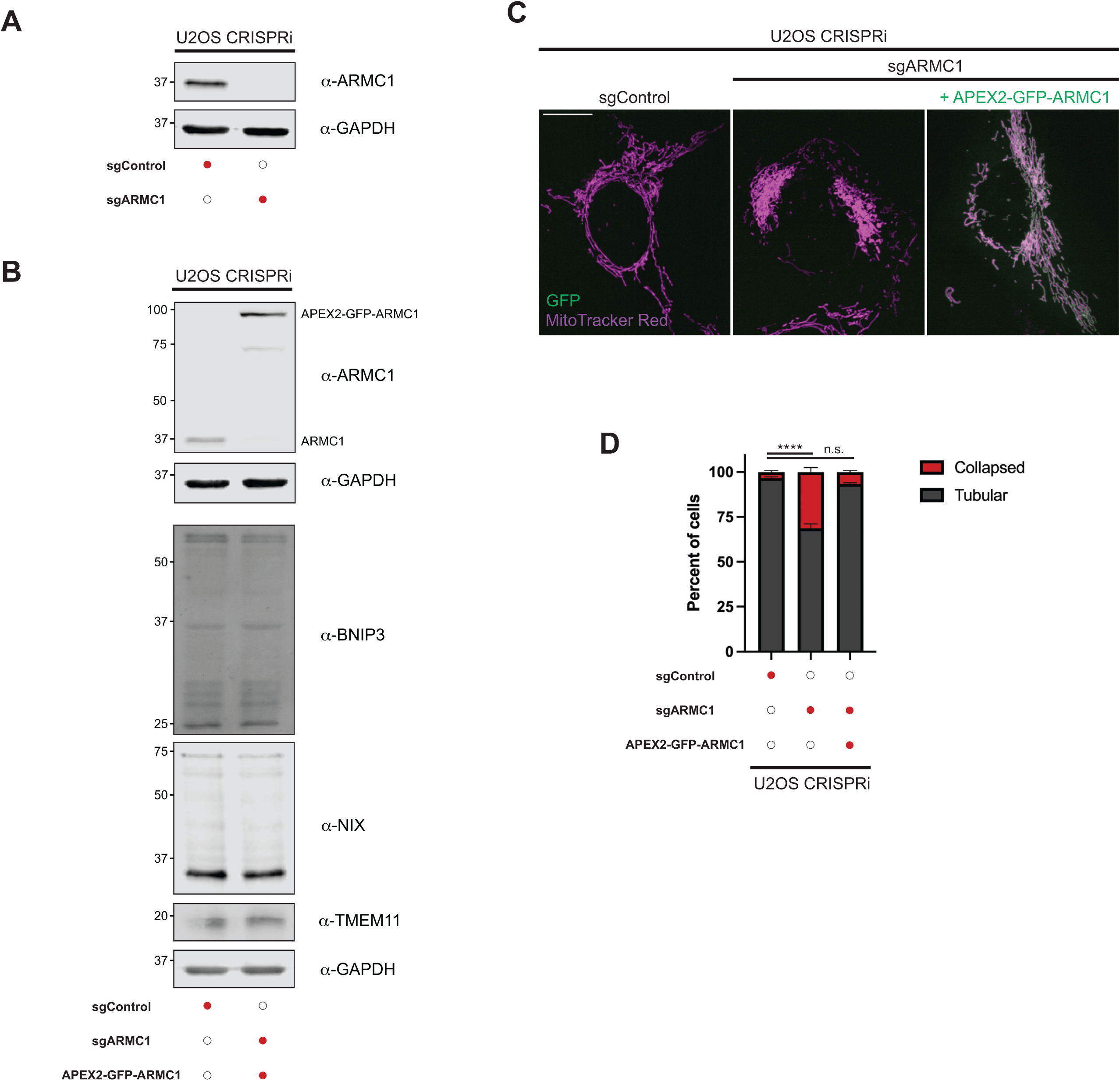
Characterization of APEX2-GFP-ARMC1-expressing cells. **(A)** Western analysis is shown with the indicated antibodies of lysate generated from U2OS CRISPRi cells expressing control or ARMC1 sgRNA. **(B)** Western analysis is shown with the indicated antibodies of lysate from U2OS control CRISPRi cells or U2OS ARMC1 CRISPRi cells stably expressing APEX2-GFP-ARMC1. **(C)** Maximum intensity projection confocal images are shown of cells as in (B) stained with MitoTracker Red CMXRos. **(D)** A graph is shown of the categorization of mitochondrial morphology from cells as in (C). Data represent 50 cells per condition in each of three independent experiments and bars indicate S.E.M.. Asterisks (****p<0.0001) represent one-way ANOVA with Tukey’s multiple comparisons test of tubular morphology. N.S. indicates not statistically significant. Scale bar = 15 µm.

**Figure S2.**
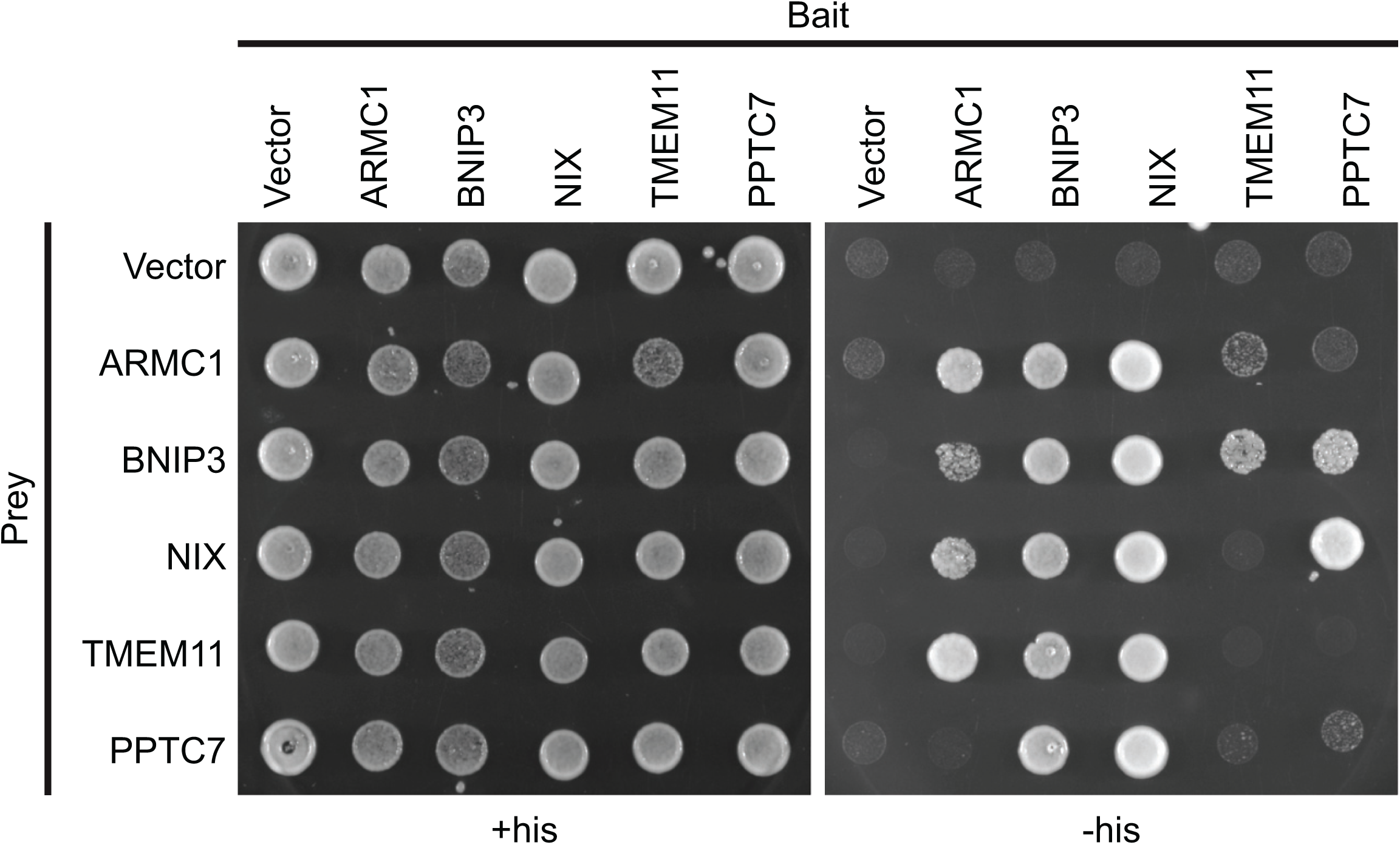
ARMC1 directly interacts with TMEM11, BNIP3, and NIX by yeast two-hybrid analysis. Images are shown of growth analysis of yeast strains expressing the indicated bait and prey constructs and spotted on permissive (+his, left) or selective (-his, right) plates.

**Figure S3.**
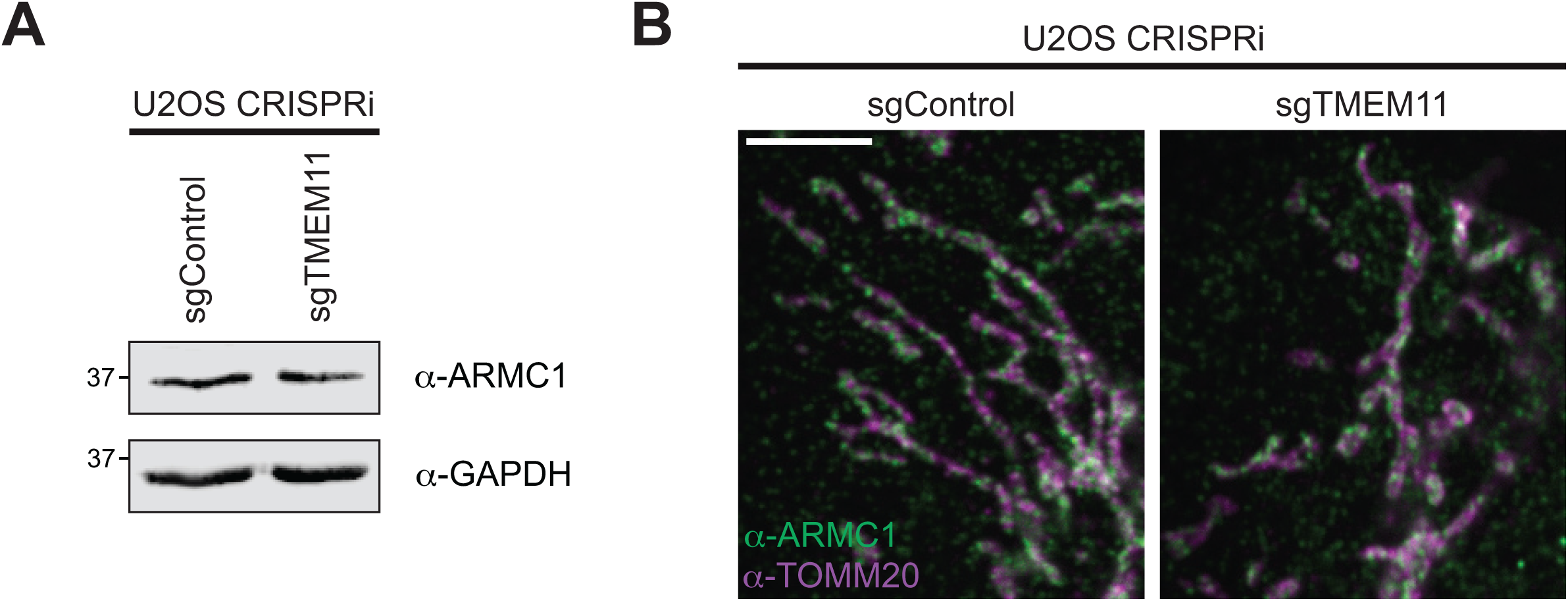
TMEM11 depletion does not affect ARMC1 levels or mitochondrial targeting. **(A)** Western analysis is shown with the indicated antibodies of lysate from U2OS CRISPRi cells expressing control or TMEM11 sgRNAs. **(B)** Single plane confocal images are shown for cells as in (A) that were fixed and stained with ARMC1 (green) and TOMM20 (magenta) antibodies. Scale bar = 5 µm.

**Figure S4.**
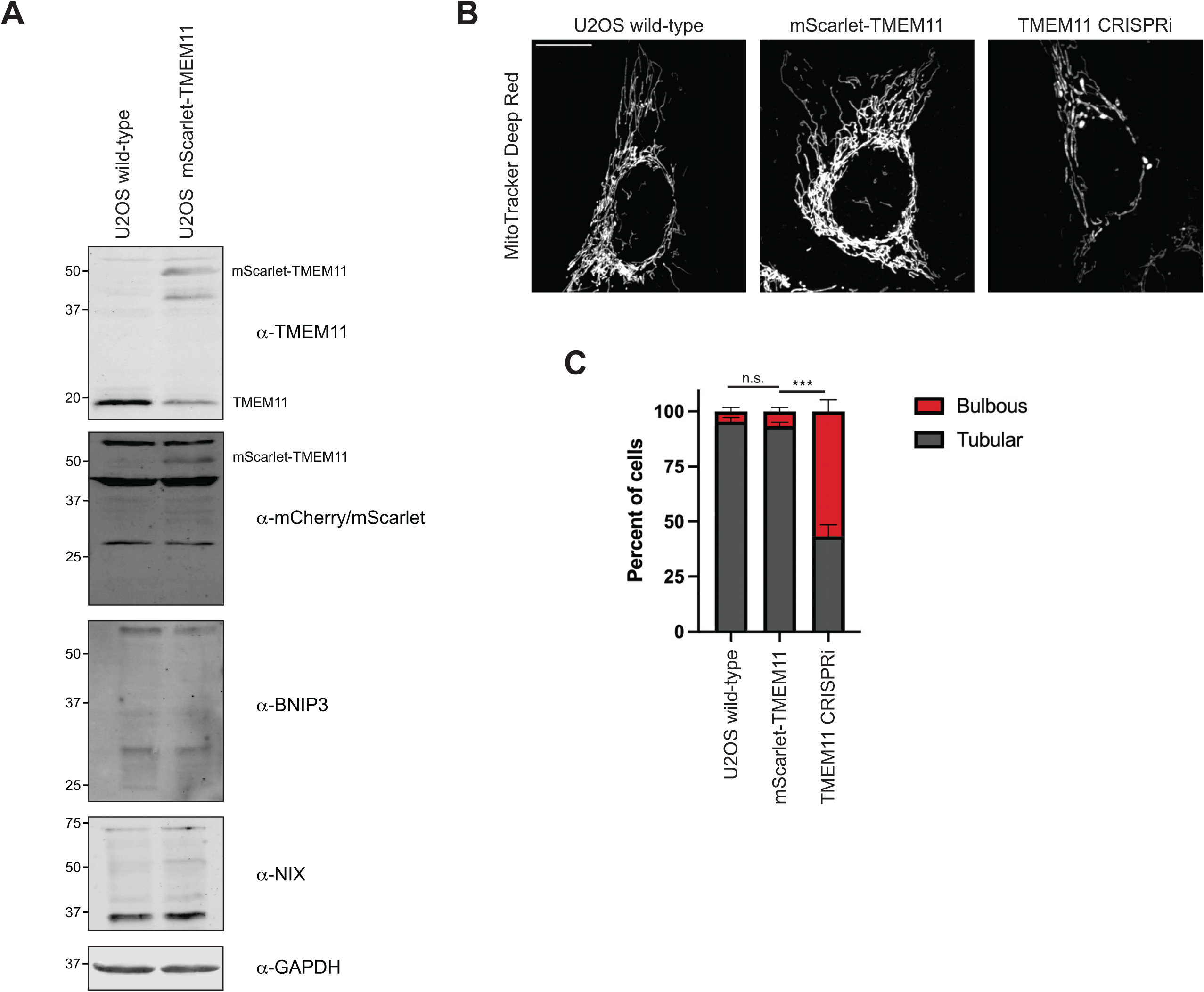
Characterization of mScarlet-TMEM11 CRISPR knock-in cells. **(A)** Western analysis is shown with the indicated antibodies of lysate from wild-type U2OS cells or mScarlet-TMEM11 CRISPR knock-in cells. **(B)** Maximum intensity projection confocal images are shown of cells as in (A), as well as U2OS TMEM11 CRISPRi cells, stained with MitoTracker Deep Red. **(C)** A graph is shown of the categorization of mitochondrial morphology from cells as in (B). Data represent 50 cells per condition in each of three independent experiments and bars indicate S.E.M.. Asterisks (***p<0.001) represent one-way ANOVA with Tukey’s multiple comparisons test of tubular morphology. N.S. indicates not statistically significant. Scale bar = 15 µm.

**Figure S5.**
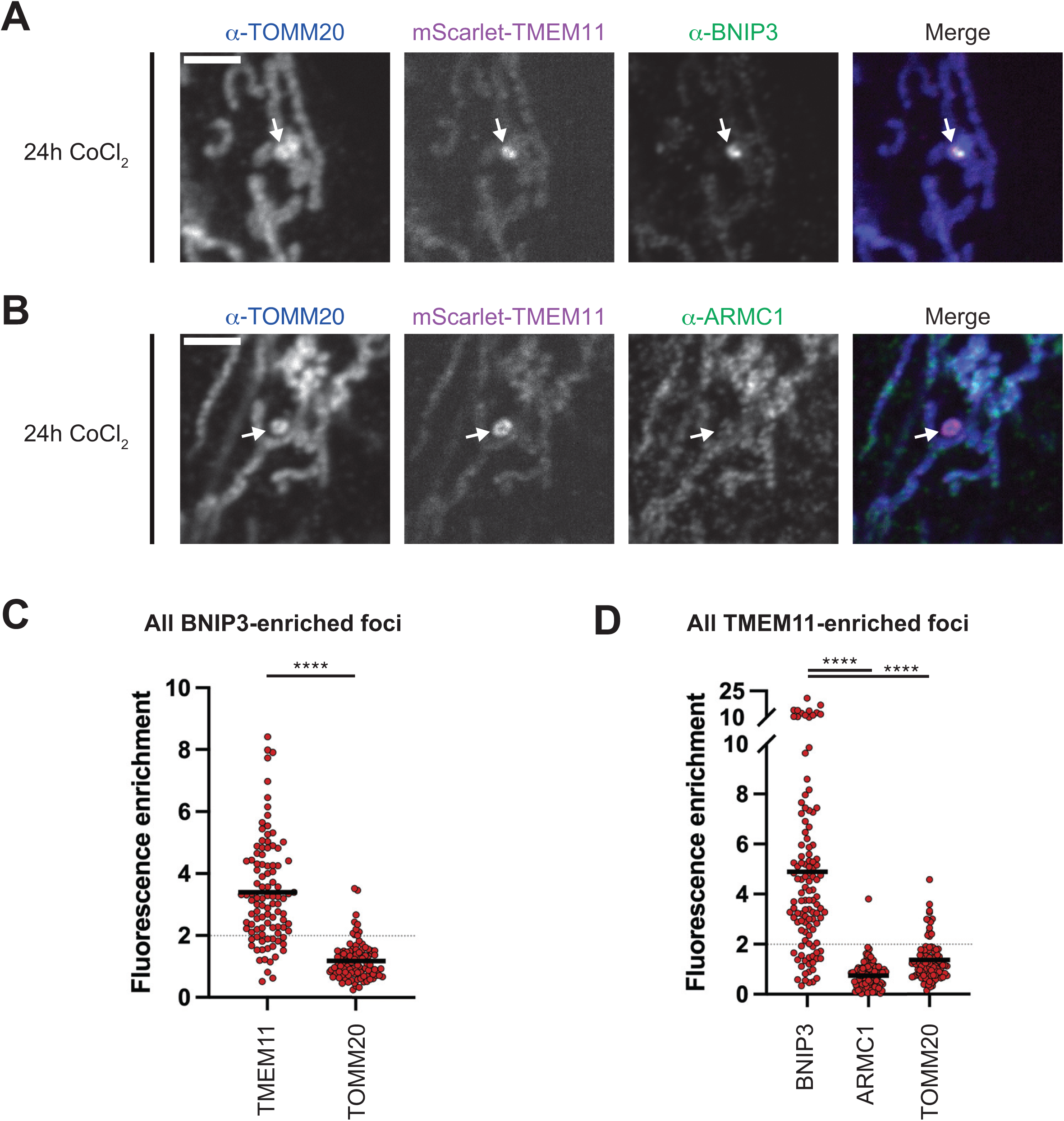
ARMC1 does not enrich at BNIP3-marked mitophagosomes formed during CoCl_2_ treatment. **(A-B)** Single plane confocal images are shown of U2OS mScarlet-TMEM11 CRISPR knock-in cells treated with CoCl_2_ for 24h and fixed and stained with the indicated antibodies. Arrows mark BNIP3 and/or TMEM11-enriched mitophagosomes. **(C)** A graph is shown of the relative fluorescence enrichment of TMEM11 or TOMM20 at BNIP3-enriched foci relative to signal at nearby mitochondrial tubules from images as in (A). Data represent n=101 foci collected in three independent experiments. **(D)** A graph is shown as in (C) of the relative fluorescence enrichment of BNIP3, ARMC1, and TOMM20 at all mScarlet-TMEM11 enriched foci from images as in (A-B). Data represent n=114 foci (BNIP3) and n=101 foci (ARMC1, TOMM20) collected in three independent experiments. Solid horizontal bars on graphs indicate means. Asterisks (****p<0.0001) represent a paired t-test (C) or one-way ANOVA with Tukey’s multiple comparisons test (D). Scale bars = 3µm.

**Figure S6.**
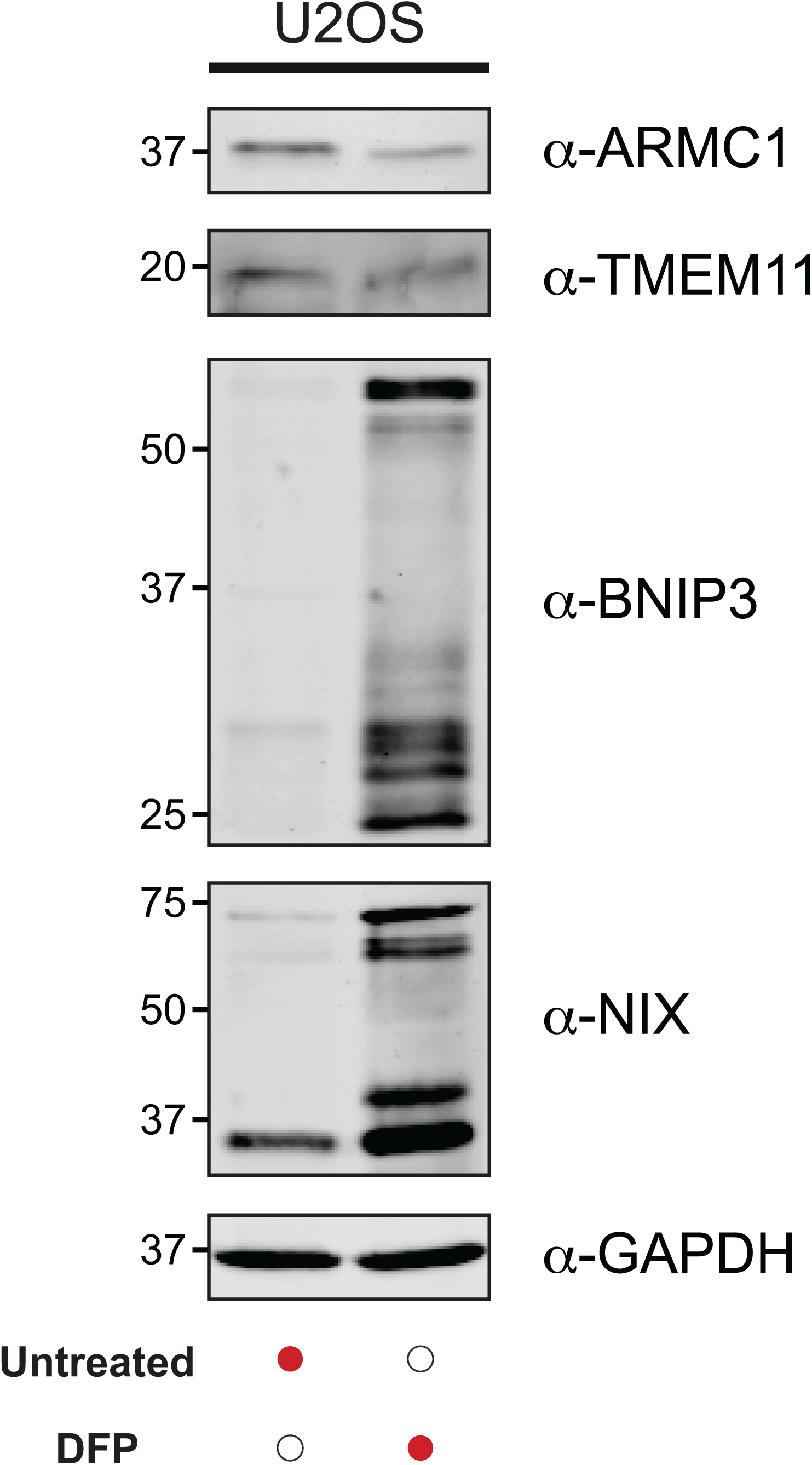
ARMC1 and TMEM11 levels do not increase after DFP treatment. Western analysis is shown with the indicated antibodies of lysate from wild-type U2OS cells before and after 1mM DFP treatment for 24h.

**Figure S7.**
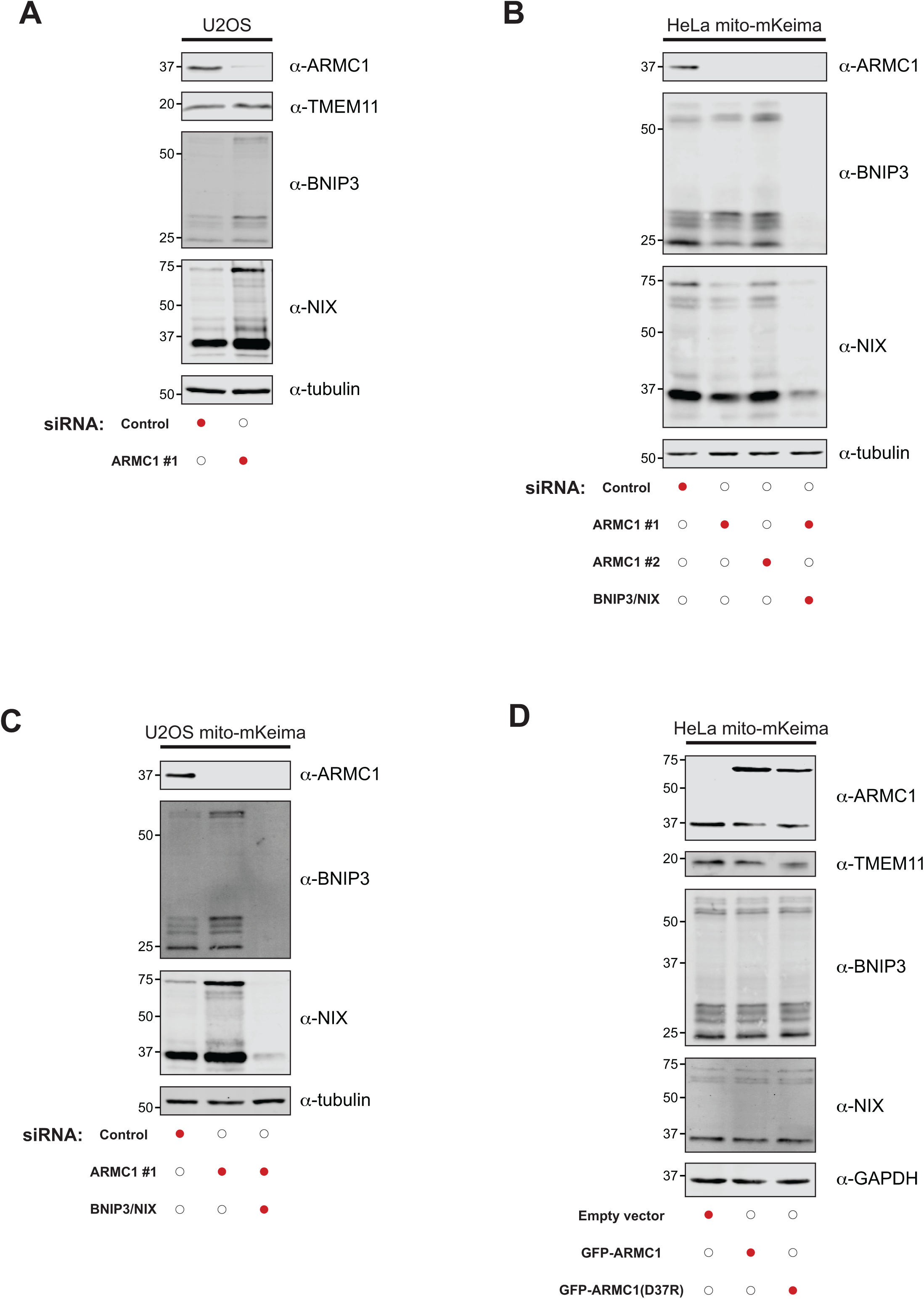
Characterization of ARMC1-depleted and -overexpressing cells. **(A)** Western analysis is shown with the indicated antibodies of lysate from wild-type U2OS cells transfected with the indicated siRNAs. **(B-C)** As in (A) for HeLa (B) or U2OS (C) cells stably expressing mito-mKeima. **(D)** Western analysis is shown with the indicated antibodies of lysate from HeLa mito-mKeima cells transfected with the indicated constructs.

**Figure S8.**
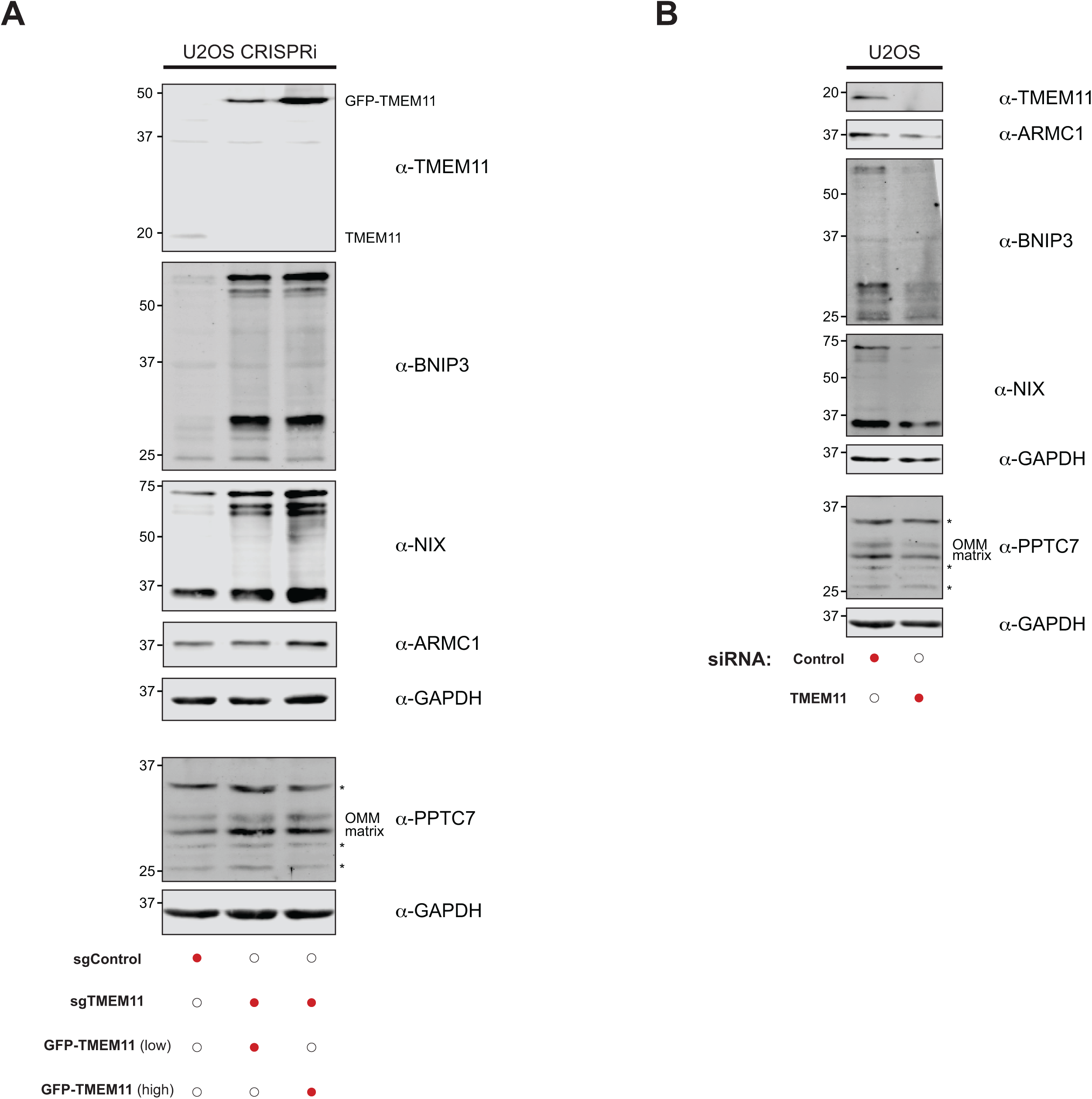
Characterization of TMEM11-depleted and -overexpressing cells. **(A)** Western analysis is shown with the indicated antibodies of lysate from U2OS control CRISPRi cells or TMEM11 CRISPRi cells stably expressing GFP-TMEM11. Cells were sorted for low or high expression levels of GFP-TMEM11 by FACS (*13*). PPTC7 blots are annotated for the OMM and matrix forms of the protein. Asterisks indicate non-specific bands. **(B)** As in (A) for wild-type U2OS cells transiently transfected with the indicated siRNAs.

**Figure S9.**
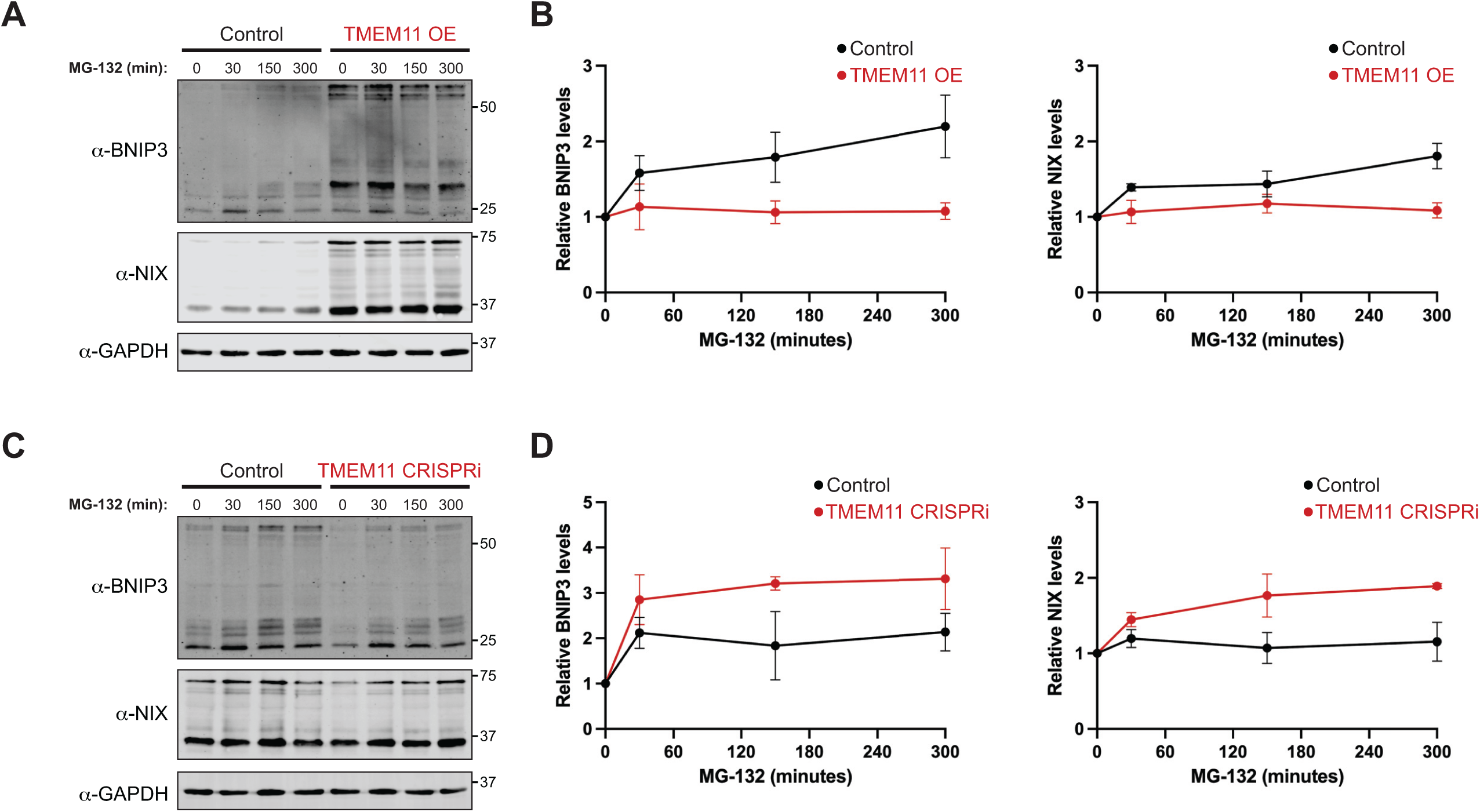
TMEM11 promotes stabilization of BNIP3 and NIX. **(A)** Representative western blots are shown with the indicated antibodies of lysates from U2OS control CRISPRi cells (left) or TMEM11 CRISPRi cells stably expressing high levels of GFP-TMEM11 (right). Cells were treated with 10 µM MG-132 for the indicated times. **(B)** Graphs are shown of quantifications of the relative levels of BNIP3 (left) or NIX (right) after treatment with MG-132 as in (A). The BNIP3 levels for control (black lines) and GFP-TMEM11 overexpressing cells (red lines) were normalized to GAPDH signal and determined relative to the 0-minute timepoint. Data shown are from three independent experiments and bars indicate S.E.M.. **(C-D)** As in (A-B) for U2OS CRISPRi control cells (black lines) and U2OS TMEM11 CRISPRi cells (red lines).

## Materials and Methods

### Cell culture

HeLa, HeLa mito-mKeima (*41*), HEK 293T, U2OS, and U2OS CRISPRi (*42*) cells were generous gifts of Mariusz Karbowski, Richard Youle, and Jodi Nunnari. U2OS mito-mKeima cells and U2OS TMEM11 CRISPRi cells with or without stable expression of APEX2-GFP-TMEM11 or GFP-TMEM11 were previously described (*13*). All cell lines were cultured in DMEM (D5796; MilliporeSigma) supplemented with 10% fetal bovine serum (F0926; MilliporeSigma), 25 mM HEPES (H0887; MilliporeSigma), and 1% penicillin/streptomycin (P4333; MilliporeSigma). For all experiments, cells were maintained at low passage number and regularly tested for mycoplasma contamination. All pseudohypoxia treatments were performed with deferiprone (DFP) (379409; MilliporeSigma) or CoCl_2_ (C8661; MilliporeSigma) for 24 hours.

### Plasmids and siRNA oligonucleotides

For transient transfections, pGFP-TMEM11, pmCherry-LC3, and pcDNA3.1-PPTC7-GFP were described previously (*13, 34*). pGFP-ARMC1 was generated by PCR amplifying ARMC1 from cDNA generated from U2OS cells and cloning into the XhoI/BamHI sites of pAcGFP1-C1 (Takara Bio) by Gibson assembly. pGFP-ARMC1(D37) was subsequently generated by PCR amplification and Gibson assembly. pGFP-FIS1(122–152) (GFP-OMM) was generated by PCR amplification of GFP with primers containing the coding sequence for human FIS1(amino acids 122-152) and cloning into the XhoI/BamHI sites of pAcGFP1-C1 by Gibson assembly. pmCherry-TMEM11 was generated by subcloning the TMEM11 coding cassette from pGFP-TMEM11 into the XhoI/BamHI sites of pmCherry-C1 (Takara Bio). For yeast two-hybrid analysis, pGADT7-BNIP3, pGBKT7-BNIP3, pGADT7-NIX, pGBKT7-NIX, pGADT7-TMEM11, pGBKT7-TMEM11, pGADT7-PPTC7 and pGBKT7-PPTC7 were described previously (*13, 18*). pGADT7-ARMC1 and pGBKT7-ARMC1 were generated by PCR amplifying the ARMC1 coding cassette from pGFP-ARMC1 and cloning into the NdeI/BamHI sites of pGADT7 and pGBKT7 (Takara Bio), respectively, by Gibson assembly.

For CRISPRi-mediated stable knockdown, ARMC1 sgRNA-expressing plasmid was generated by annealing the following forward and reverse oligonucleotides into pU6-sgRNA EF1alpha-puro-T2A-BFP (*43*) digested with BstXI and BlpI: Forward: 5’-TTGGTCAGGGAGCCAAAGAGCGAGTTTAAGAGC-3’; Reverse: 5’-TTAGCTCTTAAACTCGCTCTTTGGCTCCCTGACCAACAAG-3’. To stably express APEX2-GFP- ARMC1, PCR-amplified APEX2 and GFP-ARMC1 was cloned into XhoI/BamHI-digested pLVX-puro (Takara Bio) using Gibson Assembly.

For siRNA-mediated knockdown, Silencer Select siRNAs (Negative control no. 2 (4390846); ARMC1 (#1; s30304); ARMC1 (#2; s30305); TMEM11 (s16855); BNIP3 (s2060); NIX (s2063); Thermo Fisher Scientific) were used. Except where indicated, ARMC1 siRNA #1 was used in all experiments. Sequences of siRNAs are as follows:

ARMC1 (#1): 5′-GCAAGUUGUGAAAAGUGAATT-3′,

ARMC1 (#2): 5′- GCAGAUCCGUUAAACAGAATT-3′,

TMEM11: 5′-GAUUAUUCCCACUACAUUUTT-3′

BNIP3: 5′-CCCAUAGCAUUGGAGAGAATT-3′

NIX: 5′-GAUUCUUUUGGAUGCACAATT-3′

### Transient transfections

Approximately 150,000 cells were seeded per well of a 6-well cell culture plate ∼12 h prior to transfection. For transient expression in wild-type contexts, plasmid transfections were performed using Lipofectamine 3000 for 5–6 h and then cells were passaged to glass-bottom imaging or culture dishes for subsequent experiments. For transient knockdowns, transfections were performed using Lipofectamine RNAiMAX (Thermo Fisher Scientific) in 6-well cell culture plates with the indicated siRNAs (20 nM). Cells were incubated with the liposome/siRNA mixture for 24h, followed by passage of approximately 150,000 cells to new 6-well culture plates. Cells were allowed to adhere for ∼12 h and a second round of transfection was performed using Lipofectamine 3000 for 5-6 h with the indicated siRNA (20 nM) and any additional plasmids, where indicated. After a second round of transfection, cells were passaged to glass-bottom imaging or culture dishes for subsequent experiments.

### Stable cell line generation

Lentiviral plasmids were transiently transfected into HEK 293T cells with standard packaging plasmids, and supernatant was harvested and filtered through a 0.45 µm PES filter.

To generate U2OS ARMC1 CRISPRi cells, U2OS cells stably expressing the dCas9-KRAB transcriptional repressor (*42*) were transduced with ARMC1 sgRNA viral supernatant and 5.33 µg/ml polybrene (TR-1003-G; MilliporeSigma), and the top 50% of BFP-expressing cells were sorted by FACS in the UTSW Flow Cytometry Core Facility. U2OS ARMC1 CRISPRi cells were subsequently transduced with APEX2-GFP-ARMC1, and the lowest 25% of GFP-expressing cells were sorted by FACS.

Chromosomal knock-in of mScarlet upstream of exon 1 of *TMEM11* in U2OS cells was performed using homology-directed repair essentially as described previously (*44*). Briefly, TMEM11 sgRNA-expressing plasmid was generated by annealing the following forward and reverse primers, designed with CRISPOR (*45*), into pX459V2.0-HypaCas9 ((*46*); Addgene 108294) that was digested with BbsI and phosphatase-treated: 5’-CACCGCCAACGTCTCGCGCCAAGG-3’ and 5’-AAACCCTTGGCGCGAGACGTTGGC-3’. Homology arms with approximately 800 base pairs upstream and downstream of the TMEM11 start codon, with silent mutations corresponding to potential PAM sites to avoid cutting by Cas9, were synthesized by Twist Biosciences. These homology arms were cloned, flanking the mScarlet coding sequence along with DNA coding for the synthetic linker GSGSGSGSGSGS, into an empty vector backbone using Gibson assembly. These plasmids were co-transfected into U2OS cells using Lipofectamine 3000 (Thermo Fisher Scientific), incubated for 24h with 3 µg/ml puromycin (Thermo Fisher Scientific) to enrich for transfected cells, allowed to recover, and finally FACS sorted to enrich for a population of majority (∼90%) mScarlet-expressing cells.

### Mitochondrial isolation

Mitochondria were isolated by differential centrifugation as described previously (*13*). The indicated cells (U2OS and derivatives) were grown to approximately 90% confluency in 15 cm culture dishes, washed once with pre-warmed DPBS (MilliporeSigma D8537), harvested by scraping into pre-warmed DPBS, and collected by centrifugation (200 × g, 5 min). Cells were resuspended in approximately 5–10 pellet volumes of ice-cold mitochondria isolation buffer (10 mM Tris/MOPS pH 7.4, 0.25 M sucrose, 1 mM EGTA) and homogenized with 25-30 strokes of a glass dounce. The lysate was centrifuged at low speed (600 × g, 10 min, 4°C) to remove unbroken cells and nuclei. Crude mitochondria were enriched by centrifugation (10000 × g, 15 min, 4°C) and the mitochondrial pellet was resuspended in cold mitochondria isolation buffer. The concentration of crude mitochondria was determined by a Bradford assay (BioRad) and 100 µg aliquots were flash frozen in liquid nitrogen and stored at −80°C. For pseudohypoxia treatment (Fig. 2E), U2OS cells were treated with 300 μM DFP for 24 h and mitochondria were isolated as described above. For siRNA-mediated knockdown of ARMC1 (Fig. 2F), mitochondria were isolated from U2OS cells that were subjected to two rounds of transfection with the indicated siRNAs as described above.

### Whole-cell lysate preparation and western analysis

Whole-cell lysates were prepared by harvesting trypsinized cells, washing once with pre-warmed DPBS, and lysing in 1× RIPA buffer (50 mM Tris HCl, pH 7.5, 150 mM NaCl, 1% sodium deoxycholate, 0.1% SDS, 1% NP-40, 1 mM EDTA) supplemented with 1× protease inhibitor cocktail (539131; MilliporeSigma). Protein concentrations were determined using a BCA assay and normalized before adding 6× Laemmli buffer (6% SDS (w/v), 21.6% glycerol (v/v), 0.18 M Tris HCl pH 6.8, 0.01% bromophenol blue (w/v), 10% β-mercaptoethanol (v/v)) to a final concentration of 1×. Samples were heated for 5 min at 95°C before equivalent amounts of lysate were resolved on Tris-Glycine polyacrylamide gels. Following electrophoresis, proteins were transferred to PVDF membranes (0.45 µm pore size) and immunoblotted with the following primary antibodies: rabbit anti-ARMC1 (HPA026085; MilliporeSigma) rabbit anti-BNIP3 (44060; Cell Signaling Technology), rabbit anti-NIX (12396; Cell Signaling Technology), rabbit anti-TMEM11 (16564-1-AP; Proteintech; or HPA062854; MilliporeSigma), rabbit anti-PPTC7 (NBP1-90654; Novus Biologicals), rabbit anti-VDAC1/2 (10866-1-AP; Proteintech), mouse anti-TOMM20 (56783; Abcam), mouse anti-mCherry (PA5-34974; Thermo Fisher Scientific), mouse anti-GAPDH (60004-1-Ig; Proteintech), mouse anti-alpha Tubulin (66031-1-Ig; Proteintech). The following secondary antibodies were used for detection: anti-rabbit DyLight 800 (SA5-35571; Thermo Fisher Scientific), goat anti-mouse DyLight 800 (SA5-35521; Thermo Fisher Scientific), or goat anti-mouse DyLight 680 (35518; Thermo Fisher Scientific). The images were acquired with an Odyssey Infrared Imaging System (LI-COR). Linear adjustments to images were made and quantification of unprocessed images was performed using Image Studio Lite (LI-COR).

### Immunoprecipitations

The relevant cell lines were cultured in 15 cm dishes to approximately 90% confluency, trypsinized, and harvested by centrifugation (300 × g, 5 min). Cells were washed once with pre-warmed DPBS and crosslinked for 30 min at room temperature with 1mM DSP (Thermo Fisher Scientific). Crosslinking was quenched by the addition of 100 mM Tris pH 7.5, and cells were collected using centrifugation (300 × g, 5 min). Cells were then lysed on ice for 30 min in three pellet volumes of immunoprecipitation lysis buffer (IPLB) (20 mM HEPES-KOH, pH 7.4, 150 mM KOAc, 2 mM Mg(Ac)_2_, 1 mM EGTA, 0.6 M sorbitol) supplemented with 1% digitonin (MilliporeSigma) and 1× protease inhibitor cocktail. The cell lysate was then centrifuged (11,500 × g, 10 min, 4°C), supernatant was collected, and protein concentration was estimated with a BCA assay. Equivalent amounts of cell lysate (1.25 mg) were mock-treated or incubated with 5 μg of anti-GFP antibody (ab290, AbCam) for 4h at 4°C with gentle agitation. Cell lysate was subsequently incubated with 12.5 µl of μMACS protein G beads (Miltenyi) for 4h at 4°C with gentle agitation to capture antibodies. Beads were then isolated from the lysate using a μ column and μMACS separator (Miltenyi), washed three times with 800 µl of IPLB supplemented with 0.1% w/v digitonin and 1× protease inhibitor cocktail, and two times with 500 µl of IPLB. Proteins bound to the beads were then eluted with 2 × 25 µl of 2× Laemmli buffer pre-warmed to 95°C, followed by western analysis as described above.

### 2D BN-PAGE analysis

Aliquots of isolated mitochondria (100 μg) were thawed on ice and centrifuged (21,000 × g, 10 min, 4°C) to remove residual buffer. Mitochondrial pellets were resuspended in 20 μl of 1× NativePAGE Sample Buffer (Thermo Fisher Scientific) supplemented with 1× protease inhibitor cocktail and digitonin to a final detergent:protein ratio of 6 g/g. Samples were solubilized on ice for 15 min, followed by centrifugation (21,000 × g, 30 min, 4°C). Coomassie Blue G-250 dye was added to the supernatant to a final detergent:dye ratio of 16 g/g. Samples were then run under native conditions on a 3-12% NativePAGE Mini Protein Gel (Thermo Fisher Scientific) as per the manufacturer’s instructions. Gel lanes were then manually excised and incubated in 10 ml of denaturing buffer (0.12 M Tris-HCl, pH 6.8, 4% SDS, 20% glycerol, and 10% β-mercaptoethanol) for 25 min, microwaved for 10 s halfway through incubation (*47*). Gel slices were then loaded horizontally on a denaturing Tris-Glycine polyacrylamide gel and sealed in position by polymerizing with 0.75% agarose dissolved in SDS-PAGE running buffer, followed by western analysis as described above. Protein complex sizes were determined with NativeMark Unstained Protein Standard (Thermo Fisher Scientific) which was Coomassie-stained after native gel separation and manually aligned to western blots post-processing.

### Yeast two-hybrid analysis

Yeast two-hybrid analysis was performed as described previously (*13*) with the Matchmaker Gold Yeast Two-Hybrid System (Takara Bio). Y2H Gold and Y187 yeast strains were transformed by lithium acetate transformation with bait (pGBKT7 plasmids and derivatives) and prey (pGADT7 plasmids and derivatives), respectively. Haploid strains expressing bait and prey constructs were mated on YPD plates (1% yeast extract, 2% peptone, and 2% glucose) for 24 h, followed by diploid selection on synthetic dextrose (SD; 0.7% yeast nitrogen base, 2% glucose, and amino acids) -leu-trp plates. Strains were exponentially grown in liquid SD-leu-trp media and harvested by centrifugation (6000 × g, 2 min, room temperature). Cell pellets were resuspended in water at a density of 0.5 OD_600_/ml and then spotted on SD-leu-trp (permissive) and SD-leu-trp-his (selection) plates. Plates were incubated at room temperature prior to analysis.

### Confocal fluorescence microscopy

All confocal fluorescence microscopy was carried out on a Nikon Spinning Disk Confocal microscope with Yokogawa CSU-W1 SoRa and equipped with a Hamamatsu Orca-Fusion sCMOS camera, a Nikon 100x 1.45 NA objective, and an environmental chamber. For live-cell imaging, cells were maintained in an environmental chamber at 37°C, 5% CO_2_. For all experiments, cells were grown in glass-bottom imaging dishes (CellVis). Images were acquired with Nikon Elements software using the standard spinning disk module and all z-series images were obtained with a 0.2 μm step size. ImageJ/Fiji was used for all subsequent image analysis.

### Analysis of mitochondrial morphology

Cells were treated with 25 nM MitoTracker Red CMXRos (M7512; Thermo Fisher Scientifc) (Fig. S1C) or 25 nM MitoTracker Deep Red FM (M22426; Thermo Fisher Scientific) (Fig. S4B) for ∼15 min at 37°C, washed twice with pre-warmed media, and imaged live. To quantify mitochondrial morphology, images were blinded to sample identity and manually categorized as described in the figure legends.

### Mito-mKeima mitophagy analysis

siRNA-mediated knockdown or transient transfection of the indicated plasmids in HeLa mito-mKeima or U2OS mito-mKeima cells was performed as described above prior to imaging. Neutral and acidified mito-mKeima signals were imaged using laser excitation at 488 nm and 561 nm, respectively, and emission was detected between 573 and 648 nm. In GFP co-expressing cells, GFP signal was imaged using laser excitation at 488 nm, and emission was detected between 515 and 555 nm. The identity of samples was blinded prior to analysis and the number of acidified mito-mKeima puncta per cell was manually counted in ImageJ/Fiji by assessing single plane images throughout the z-series for individual cells.

### Immunofluorescence staining

Cells cultured on glass-bottom cover dishes were fixed in 4% paraformaldehyde in PBS (Thermo Fisher Scientific) (15 min, room temperature). Fixed cells were permeabilized with 0.1% Triton X-100 (Thermo Fisher Scientific) in PBS (5 min, room temperature), rinsed with PBS, and blocked with PBS supplemented with 0.1% Triton X-100 and 10% FBS (30 min at room temperature). Cells were then incubated with the following primary antibodies, where indicated, in blocking buffer overnight at 4°C: rabbit anti-ARMC1 (HPA026085; MilliporeSigma), rabbit anti-BNIP3 (44060; Cell Signaling Technology), and/or mouse anti-TOMM20 (56783; Abcam). Cells were washed several times with PBS and subsequently incubated with one or more of the following appropriate secondary antibodies in blocking solution for 30 min at room temperature: donkey anti-mouse Alexa 405 plus (A48257; Thermo Fisher Scientific) donkey anti-rabbit Alexa Fluor 488 (A-21206; Thermo Fisher Scientific), donkey anti-mouse Alexa Fluor 555 (A-31570; Thermo Fisher Scientific), donkey anti-rabbit Alexa Fluor 555 plus(A-32794; Thermo Fisher Scientific), donkey anti-mouse Alexa Fluor 647 (A-31571; Thermo Fisher Scientific), or donkey anti-rabbit Alexa Fluor 647 Plus (A32795; Thermo Fisher Scientific). Cells were washed several times with PBS prior to imaging.

### Fluorescence enrichment determination at BNIP3- or TMEM11-marked mitophagy sites

For analysis of mScarlet-TMEM11 co-enrichment with BNIP3 after DFP (Fig. 2C) or CoCl_2_ (Fig. S5C) treatment, visually apparent and enlarged BNIP3-enriched sites on, or immediately adjacent to, mitochondrial tubules (marked with TOMM20) were identified blind to mScarlet-TMEM11 signal. Regions of interest (ROIs) were defined at the BNIP3-enriched site (ROI_foci_) and on a corresponding nearby well-resolved mitochondria (ROI_mito_) in maximum intensity projections. The fluorescence enrichment of the maximum BNIP3 signal at ROI_foci_ relative to ROI_mito_ was determined after background signal subtraction. This enrichment was found to be greater than 2-fold at 99% of analyzed sites, and thus a 2-fold cutoff was chosen for all subsequent analysis. The ROI_foci_ and ROI_mito_ values were similarly determined for mScarlet-TMEM11 and TOMM20 signal at each BNIP3-enriched site and their relative enrichments were plotted. To compare the relative enrichment of PPTC7-GFP at BNIP3-enriched sites in the presence or absence of TMEM11 (Fig. 6D), a similar analysis was performed. To determine enrichment of BNIP3, ARMC1, and TOMM20 at mScarlet-TMEM11-enriched sites (Fig. 2D, Fig. S5D), putative mScarlet-TMEM11 enrichment sites were blindly identified as described above for BNIP3 and all 2-fold enriched ROIs were subsequently analyzed.

### Determination of relative BNIP3 levels by immunofluorescence

To determine relative levels of BNIP3 between transfection or co-transfection conditions (Fig. 4E, Fig. 4H, Fig. 6B), cells were fixed and stained for BNIP3 and TOMM20 and imaged with identical imaging settings in each channel for all samples within experiments. Cells were picked for analysis blind to BNIP3 signal and the background-subtracted maximum fluorescence intensity for BNIP3 and GFP and/or mCherry signal was determined for each cell on a region where the signal was clearly evenly distributed on mitochondrial tubules defined by TOMM20 staining. Cells were chosen for subsequent analysis where relative GFP and/or mCherry expression was within a defined narrow range that allowed for comparison between samples. To generate plots, the mean BNIP3 signal for all empty vector (Fig. 4H, Fig. 6B) or non-transfected (Fig. 4E) control cells was determined for an experimental replicate, and the BNIP3 levels relative to the control mean value for that experiment were determined for each cell. Non-transfected cells (Fig. 4E) were identified as having no discernible GFP signal in the GFP-OMM transfection condition.

### Pulse-chase analysis of BNIP3 and NIX levels

For MG-132 pulse-chase experiments, cells were treated with 10 μM MG-132 (M7449; MilliporeSigma) for 30-300 min or with 0.1% ethanol as a vehicle control. For cycloheximide pulse-chase experiments, cells were pre-treated with 1mM DFP for 24 h to increase BNIP3 and NIX levels. Subsequently, cells were treated with 50 μg/mL cycloheximide (C6798; MilliporeSigma) for 30-300 min or 0.1% ethanol as a vehicle control. Samples were collected at defined times by trypsinization and centrifugation. Western analysis was performed as described above and quantitative analysis of BNIP3 and NIX signal was performed for each sample by normalization to a GAPDH loading control and relative to the vehicle treatment using Image Studio Lite.

### Statistical analyses

All statistical analyses were performed as described in the legends with GraphPad Prism 11.

## Acknowledgements

We thank members of the Friedman Lab for helpful discussions. We thank Holly Merta for technical advice. We thank Richard Youle for generously providing HeLa mito-mKeima cells. We thank Marcel Mettlen for technical advice and the UT Southwestern Quantitative Light Microscopy Facility (supported in part by NIH P30CA142543) for support and access to the Nikon SoRa microscope (purchased with NIH 1S10OD028630-01 to Kate Luby-Phelps). We thank the UT Southwestern Flow Cytometry Core for assistance with FACS sorting. This work was supported by grants from the National Institutes of Health (R35GM137894 to J.R.F.) and the National Science Foundation (#2327631 to N.M.N.).

